# Coordinated macrophage and T cell interactions mediate response to checkpoint blockade in colorectal cancer

**DOI:** 10.1101/2025.02.12.637954

**Authors:** Guillaume Mestrallet, Matthew Brown, Natalie Vaninov, Nam Woo Cho, Leandra Velazquez, Aparna Ananthanarayanan, Matthew Spitzer, Nicolas Vabret, Cansu Cimen Bozkus, Robert M Samstein, Nina Bhardwaj

**Affiliations:** Division of Hematology and Oncology, Hess Center for Science & Medicine, Tisch Cancer Institute, Icahn School of Medicine at Mount Sinai, New York, NY, USA; The Marc and Jennifer Lipschultz Precision Immunology Institute, Department of Immunology and Immunotherapy, Department of Radiation Oncology, Icahn School of Medicine at Mount Sinai, New York, NY 10029, USA; Department of Radiation Oncology and the Department of Otolaryngology–Head and Neck Surgery, University of California at San Francisco, San Francisco, CA, USA; Department of Otolaryngology–Head and Neck Surgery and the Department of Microbiology and Immunology, University of California at San Francisco, San Francisco, CA, USA

**Keywords:** Mismatch repair deficiency, immune resistance, LAG3, CTLA-4, TREM2, PD-1

## Abstract

Mismatch repair deficiency (MMRd), either due to inherited or somatic mutation, is prevalent in colorectal cancer (CRC) and other cancers. While anti-PD-1 therapy is utilized in both local and advanced disease, up to 50% of MMRd CRC fail to respond. Using animal and human models of MMRd, we determined that interactions between MHC+ C1Q+ CXCL9+ macrophages and TCF+ BHLHE40+ PRF1+ T cell subsets are associated with control of MMRd tumor growth, during anti-PD-1 treatment. In contrast, resistance is associated with upregulation of TIM3, LAG3, TIGIT, and PD-1 expression on T cells, and infiltration of the tumor with immunosuppressive TREM2+ macrophages and monocytes. By combining anti-PD-1 with anti-LAG3/CTLA4/TREM2, up to 100% tumor eradication was achieved in MMRd CRC and remarkably, in >70% in MMRp CRC. This study identifies key T cell and macrophage subsets mediating the efficacy of immunotherapy in overcoming immune escape in both MMRd and MMRp CRC settings.

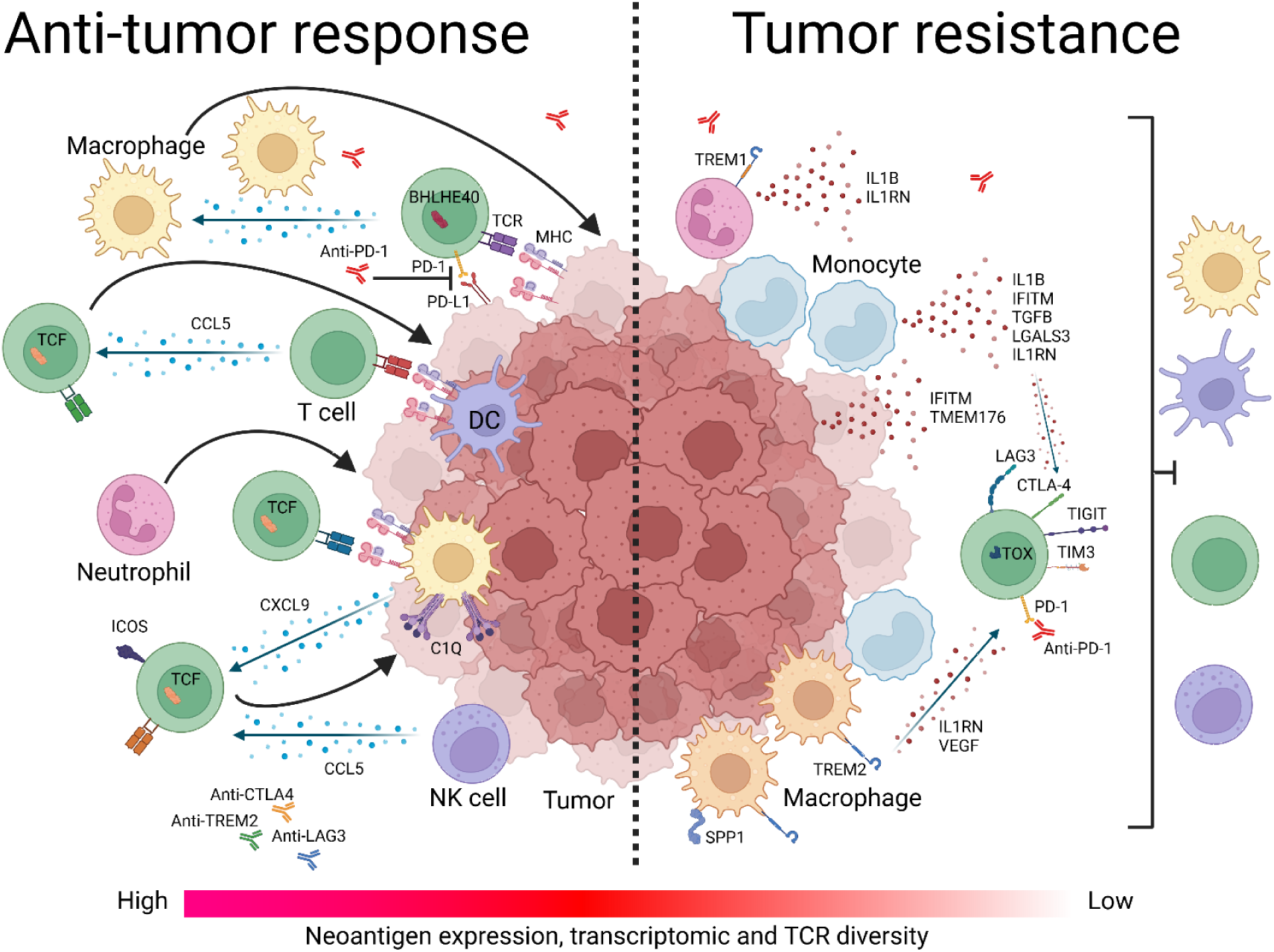

**Highlights:**

- Anti-PD-1 therapy leads to the accumulation and colocalization of MHCI/II+ C1Q+ CXCL9+ macrophages and DCs with TCF+ CCL5+ T cells that have high TCR diversity.
- Resistance to anti-PD-1 therapy involves multiple T cell checkpoints, TREM2+ macrophages, IL1B+ TREM1+ monocytes and neutrophils, and IFITM+ tumor cells.
- Simultaneous blockade of PD-1, LAG3, CTLA-4 and TREM2 dramatically prevents progression of both MMRd and MMRp tumors.
- Combination therapy completely eliminates tumors by leveraging MHC+ macrophage, CD4+ and CD8+ T cell interactions, facilitating durable anti-tumor effects.

## Introduction

Up to 30% of colorectal, endometrial, and gastric cancers exhibit deficiency in mismatch repair (MMRd) protein expression due to germline or epigenetic inactivation, leading to high genomic instability in microsatellite regions (MSI-H) ^1–5^. As a result, MMRd tumors have an increased frequency of insertions/deletions (indels) that may encode highly immunogenic, novel, shared neoantigens due to common stretches of foreign protein sequence downstream of frameshift mutations at microsatellite regions ^6^. Significantly, MMRd associated adenomas and cancers become infiltrated by T cells, contributing to the clinical efficacy of anti-PD-1 therapy ^7–10^. However, up to 50% of patients with advanced MMRd stage IV colorectal cancer (CRC) do not respond to PD-1 blockade, highlighting the need to explore resistance mechanisms impacting the success of checkpoint blockade ^11,12^.

Immune resistance can develop during tumor development, or intrinsic resistance, and after PD-1 blockade ^1^, or acquired resistance, attributed primarily to T cell exhaustion ^6,13–16^ and a dysregulated tumor microenvironment (TME) ^16,17^. In human MMRd CRC, an inflammatory hub containing T regulatory cells (Tregs), cancer-associated fibroblasts (CAFs) expressing matrix metalloproteinases (MMP), immunosupressive monocytes, and neutrophils amplify immune evasion, coupled with tumor angiogenesis and tissue remodeling ^16^. Following PD-1 blockade in MMRd CRC patients, those who achieve a complete response demonstrate an increase in CD8+ T effector memory cells, CD4+ T helper cells, and CD20+ B cells compared to patients with no complete response ^17^. This is also accompanied by a reduction in tumor infiltration by CD8+ resident memory cells, Tregs, IL1B+ monocytes, and CCL2+ fibroblasts. Other features of the TME that have been associated with resistance include the immunoediting of immunogenic neoantigens ^18^, and impaired antigen presentation in part due to the loss of beta 2 microglobulin ^1,19^. Finally, IFN-induced upregulation of HLA-E, which inhibits CD8+ T cells and NK cells via the NKG2A/CD94 receptor, has been proposed as an additional resistance mechanism ^20^.

However, the precise coordination and features of pro-immunogenic vs resistance mechanisms are still not completely defined. This affects the ability to precisely determine what pathways need to be targeted in order to overcome immune resistance in MMRd tumors and generate robust immune memory.

In this study, we employed spatial and single cell transcriptomics, spectral flow cytometry, machine learning and imaging techniques in murine tumor models along with murine and human MMRd CRC spheroid cultures to define pertinent immune resistance pathways. We uncovered evidence suggesting that targeting multiple checkpoints and macrophages overcomes resistance in MMRd tumors, and that TCF+ T cells and MHC+ macrophages orchestrate this response.

## Results

### MMRd tumors responding to anti-PD-1 are highly infiltrated by MHC+ C1Q+ macrophages

To investigate the determinants of immune response and resistance within MMRd tumors, we utilized a murine colorectal cancer CT26 cell line, in which MSH2, a DNA mismatch repair protein, was either deleted or left intact, thereby modeling MMRd or MMRp tumors, respectively. Cells were injected into the flank of BALB/C mice and after 14 days, once tumors were established, mice were treated with anti-PD-1. As expected, MSH2 KO tumor growth was substantially reduced compared to WT tumors after 28 days (**Figure 1 A**). Despite this significant reduction in MSH2 KO tumor growth following anti-PD-1 therapy, tumors persisted and grew gradually to 35 days when mice were euthanized (**Figure 1 A**). To ensure that the observed differences in tumor sizes in vivo between MSH2 KO and WT tumors were not due to different growth rates of the cell lines, we generated spheroids and followed their growth in vitro. We observed no difference in spheroid growth based on MSH2 status (**Extended Figure 1**), confirming that control of tumor growth in vivo is likely due to an induction of anti-tumor immunity. Our results confirmed that anti-PD-1 therapy contributed towards the control of MMRd tumors, as seen in human cohorts, yet therapy resistance persisted as tumors continued to grow, albeit slower.

**Figure 1.**
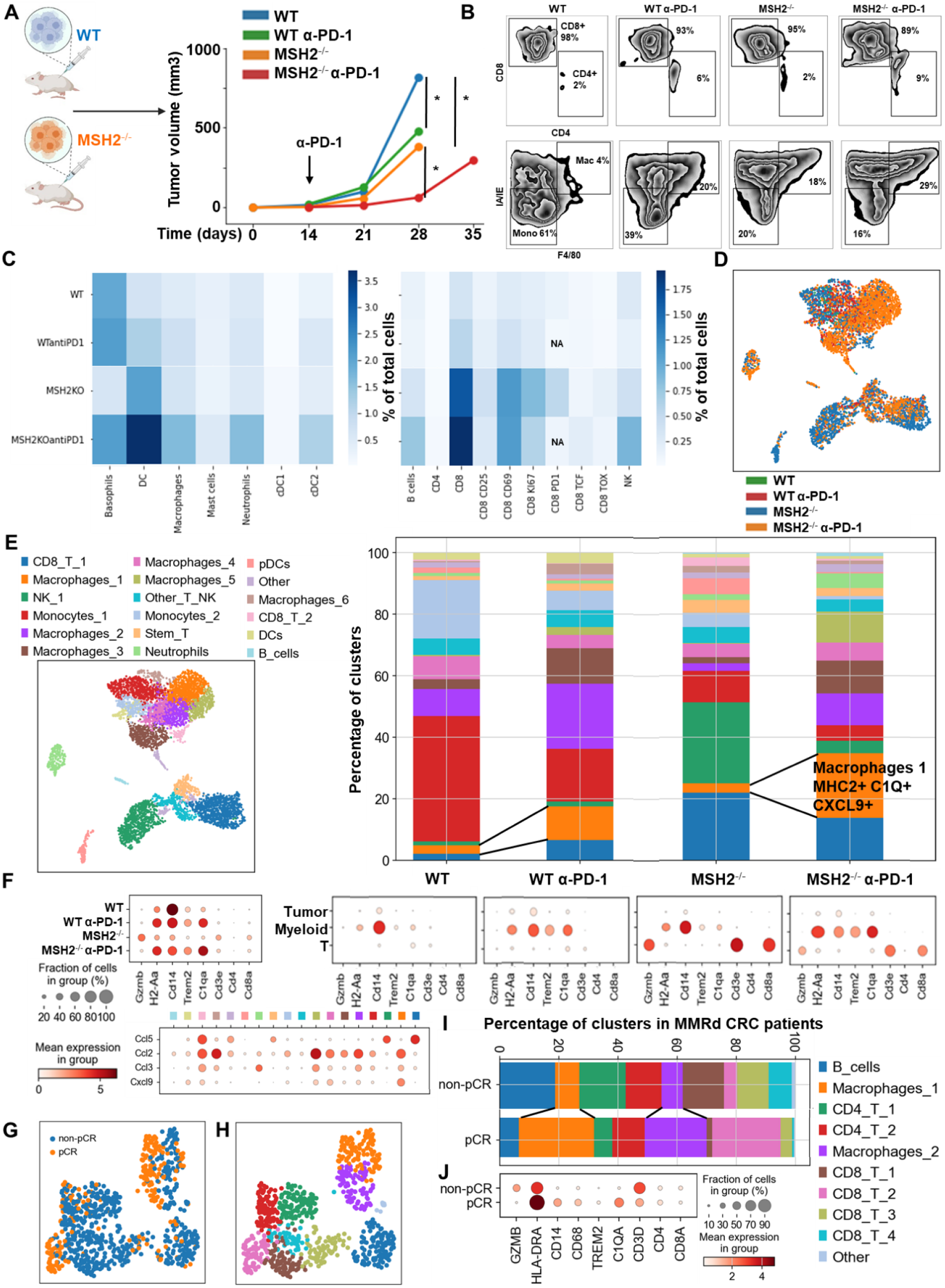
MMRd tumors responding to anti-PD-1 are highly infiltrated by MHC+ C1Q+ macrophages. **A** 200,000 CT26 tumors, either deleted or not for MSH2, were allowed to grow for 28 or 35 days in BALB/C mice, with or without anti-PD-1 administered twice a week after 14 days. **B/C** Quantification of differential lymphocyte infiltration expression according to the MSI status and ICB through spectral flow cytometry. (N = 6 replicates, Mann-Whitney U Test, p-value < 0.05). **D** ScRNAseq UMAP of immune cells in CT26 WT and MSH2 KO tumors with or without anti-PD-1 therapy. **E** UMAP of immune clusters subsequent to Leiden clustering by scanpy. Percentage of cells in each immune cluster. **F** Gene expression within each immune cluster. **G/H/I/J** We reanalyzed human MMRd CRC patient data previously published ^17^, focusing on patients who responded or did not respond to anti-PD-1 therapy alone, and the underlying T cell and macrophage profiles. **G** ScRNAseq UMAP of immune cells in patients who responded (pCR) or did not respond (non-pCR) to anti-PD-1 therapy. **H** UMAP of immune clusters subsequent to Leiden clustering by scanpy. **I** Percentage of cells in each immune cluster. **J** Gene expression within each immune cluster.

To understand which immune subsets are critical in mediating tumor growth control, we investigated immune infiltration within CT26 WT and MSH2 KO tumors, both with and without anti-PD-1 therapy. Using flow cytometry, we observed that both 14 and 28 days after inoculation, MSH2 KO tumors were more infiltrated by immune cells compared to WT tumors (12% vs 6% after 28 days, p-value < 0.05) (**Extended Figure 2 A**). Compared to WT tumors, MSH2 KO tumors had increased infiltration by DCs and CD4+ T cells at day 14, followed by enrichment of activated proliferating CD8+ T cells at day 28, including PD-1+ CD8+ T cells (**Figure 1 B/C, Extended Figure 2 A**). Interrogation of scRNAseq showed that MSH2 KO tumors also had greater numbers of natural killer (NK) cells, T cells, B cells, dendritic cells (DCs), macrophages, and neutrophils compared to the poorly infiltrated WT tumors, which showed mainly monocyte infiltration (**Figure 1 C/D/E, Extended Figure 3).** Post anti-PD-1 therapy, there was a decrease in monocytes and an increase in macrophages, T cells, CD8+ T cell subsets expressing TCF, a marker of stem-like T cells ^21^, and neutrophils, particularly notable in MSH2 KO tumors (**Figure 1 C/D/E, Extended Figure 2/3**). These macrophages expressed MHCII, C1Q, a protein that initiates the classical complement pathway of the complement system ^22^, CXCL9, a cytokine that attracts CD8+ cytotoxic T cells ^23^, but also TREM2, a marker of resistance ^24–27^ (**Figure 1 F, Extended Figure 3**). Overall, the delayed tumor growth by MSH2 KO tumors, especially following PD-1 blockade, correlates with an increased immune infiltration comprised of T cells, including activated CD8+ T cell subsets expressing TCF, DCs, neutrophils and MHC+ C1Q+ CXCL9+ macrophages.

Lastly, we compared our observations in murine models with the human condition. To do so we conducted retrospective analyses on human COAD and UCEC MMRd cohorts using CIBERSORT data available on TCGA (**Extended Figure 4 A**) and the CRI iAtlas (**Extended Figure 4 B/C**). Remarkably, the extensive MMRd tumor immune cell infiltration observed in mice (**Figure 1 C/D**) was also observed in humans (**Extended Figure 4**), with notable emphasis on Th2 and CD8+ T cells. Furthermore, we conducted a reanalysis of human MMRd CRC scRNAseq patient data, focused on patients who either achieved a complete response or did not completely respond to anti-PD-1 therapy, with examination of underlying immune profiles (**Figure 1 G/H, Extended Figure 5**) ^17^. Human MMRd CRC patients who responded to anti-PD-1 therapy had greater infiltration of macrophages expressing C1Q and MHCII (**Figure 1 I/J, Extended Figure 5**), mirroring our observations in the mouse model.

### A high diversity of CCL5+ PRF1+ T cell clones, DCs and MHC+ C1Q+ macrophages colocalize in MMRd tumors responding to anti-PD-1

We next investigated which underlying mechanisms might lead to high immune cell infiltration in MMRd tumors in their response to anti-PD-1. We explored whether this increase in immune infiltration might be attributable or correlated with an increased presence in lymph nodes and spleen. We quantified immune cell distribution in the spleen and lymph nodes 28 days after tumor injection using spectral flow cytometry. Notably, we observed a higher presence of macrophages and neutrophils in the draining lymph nodes of mice injected with MSH2 KO tumors compared to WT tumors after 28 days (**Extended Figure 2 E**). Following anti-PD-1 therapy, a surge in TCF+ CD4+ and TCF+ CD8+ T cells, DCs, neutrophils, and macrophages was also evident in the spleen of mice injected with MSH2 KO tumors after 28 days (**Extended Figure 6 C**). Moreover, there was an elevated count of TCF+ CD4+ and TCF+ CD8+ T cells in the lymph nodes **(Extended Figure 6 D**). Conversely, in mice injected with WT tumors, following anti-PD-1 therapy, a reduction in CD4+ and CD8+ T subsets was discerned in the spleen and the lymph nodes after 28 days (**Extended Figure 6 C/D**). In summary, the increased infiltration of TCF+ T cells and myeloid cells into MSH2 KO tumors following PD-1 blockade correlates with escalated presence in both spleen and lymph nodes.

We also investigated whether the high burden and the landscape of mutations in MMRd tumors shape the immune milieu within the TME leading to the elevated TCF+ T cell infiltration. To this end, we utilized different CT26 MSH2 KO cell lines, with high and low mutation loads, respectively, as determined by WES (**Extended Figure 7**). Notably, these different MSH2 KO cell lines displayed differential response to PD-1 blockade. We observed that high indel and SNV loads of the tumor before initiating ICB correlated with delayed tumor growth and an increased infiltration by TCF+ T cells (**Extended Figure 7**). Thus, high tumor mutational burden correlates with high infiltration by TCF+ T cells and response to anti-PD-1. Tumor allele frequencies of frameshift and SNV mutations also decreased following anti-PD-1 therapy, when comparing pre- and post-treatment timepoints within the same group, suggestive of immuno-editing (**Extended Figure 7**).

Next, we investigated if MSH2 KO tumors having high tumor mutational burden and consequently increased T cell infiltration were marked by changes in T cell clonal diversity. We applied scRNAseq coupled with TCR profiling to profile the diversity of tumor infiltrating T cells (**Figure 2 A**, **Extended Figure 8 A/B**). There were very few T cells in WT tumors, limiting the analysis for this condition. Only 4 clonotypes were shared between WT and MSH2 KO tumors (clones 9, 1, 19, 77), while 5 clonotypes were shared in MSH2 KO tumors with or without anti-PD-1 therapy (clones 183, 187, 125, 9, 184) (**Figure 2 B**). Notably, MSH2 KO tumors were characterized by the strong expansion of 1 clone (183) in the CD8+ PRF1+ cluster, while anti-PD-1 promoted the expansion of multiple clones (**Figure 2 B, Extended Figure 8 C/D/E**) (62 clones in MSH2 KO tumors after ICB vs 46 clones without ICB). Clone 183 was characterized by expression of the Human T Cell Receptor Alpha Variable (TRAV) genes TRAV16N and TRAV3-3 and by a higher length of CDR3 regions (**Extended Figure 8 F/G**). Interestingly, the more abundant clones in MSH2 KO tumors (183, 187) overexpress BHLHE40, a tissue-specific regulator of murine tissue-resident macrophage proliferation ^28^, repressing IL-10 and increasing IFN-γ secretion, also involved in Th1-like cell and CD8+ memory T cell responses to CD40 agonist immunotherapy in CRC ^29^. These clones also overexpressed PRF1, the gene allowing the production of Perforin and enabling cytolytic activity of the T cells, suggesting that they are able to target the tumor cells (**Figure 2 B**). The more abundant clones in MSH2 KO tumors following anti-PD-1 therapy (9, 27) also overexpress EIF3L, SERPIN and CCL5, a cytokine recruiting various immune cells such as T cells, eosinophils, basophils, monocytes, NK cells, and DCs ^30^ (**Figure 2 B**, **Extended Figure 8 H**). Overall, recognition of MSH2 KO tumors involved specific TCRs expressed by CD8+ T cells, and anti-PD-1 increased the diversity of the T cell clonal response (**Figure 2 C**), releasing CCl5 and BHLHE40 that attract other APCs and promote macrophage proliferation.

**Figure 2.**
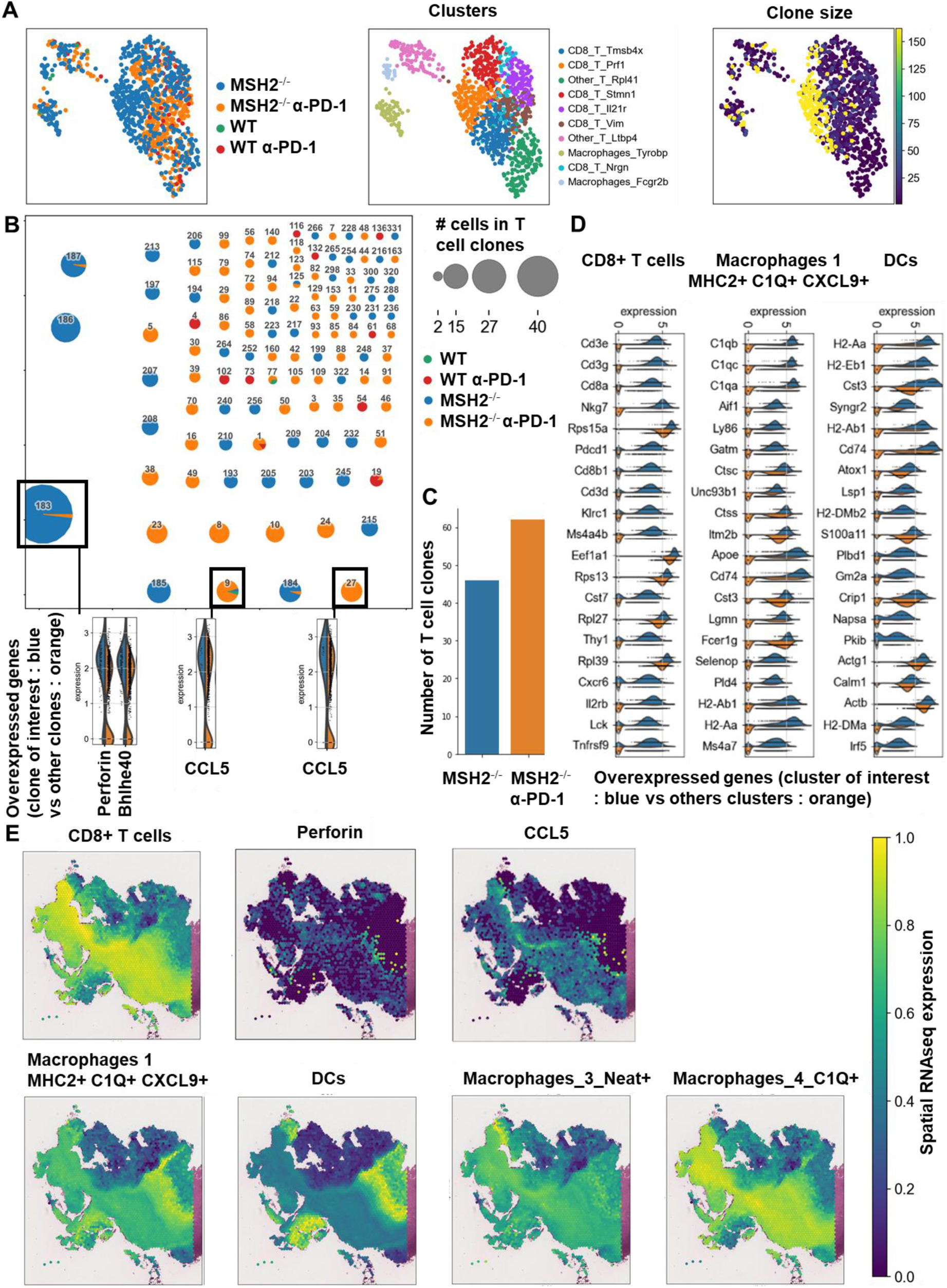
A high diversity of CCL5+ PRF1+ T cell clones, DCs and MHC+ C1Q+ macrophages colocalize in MMRd tumors responding to anti-PD-1. 200,000 CT26 tumors, either deleted or not for MSH2, were allowed to grow for 28 or 35 days in BALB/C mice, with or without anti-PD-1 administered twice a week after 14 days. **A** ScTCRseq UMAP of immune cells in CT26 WT and MSH2 KO tumors with or without anti-PD-1 therapy. UMAP of immune clusters subsequent to Leiden clustering by scanpy and clonal expansion of T cell clones. **B** Identification of T cell clones, number of cells in each clonotype for each tumor type and gene expression within each main T cell clone (clone of interest : blue vs other clones : orange). **C** Number of T cell clones according to ICB status. **D** Overexpressed genes in T cell, macrophages and DCs scRNAseq data (cluster of interest : blue vs others clusters : orange). **E** Spatial localization of each single-cell immune cluster in MSH2 KO tumors with anti-PD-1 therapy.

Subsequently, we investigated these mechanisms of cooperation between T cells and myeloid cells in the TME in situ using spatial transcriptomics. Through the application of scRNAseq clustering to spatial tumor samples, we observed that CD8+ T cells, along with macrophage clusters 3 and 4 expressing C1Q and MHCII, co-localize within the tumors (**Figure 2 D/E, Extended Figure 9**). These clusters are in proximity to DCs and macrophage cluster 1, which display high levels of C1Q and MHC expression (**Figure 2 D/E, Extended Figure 9**). The main TCR clones, overexpressing PRF1, appeared to be enriched between the T cell enriched and the T cell excluded zones, suggesting that they drive the anti-tumor response (**Extended Figure 8 J**). CD8+ T cells and NK expressed high levels of CCL5, a cytokine that attracts T cells, while the macrophage cluster 1 expressed high levels of CXCL9, a cytokine that attracts CD8+ cytotoxic T cells, and MHC molecules (**Figure 1 F**). Collectively, the infiltration of MSH2 KO cells by CD8+ T cells and macrophages expressing C1Q and MHC may account for their superior response to anti-PD-1 therapy in comparison to WT tumors. Thus, it is plausible that collaboration between DCs, macrophages and CD8+ T cells play a pivotal role in the anti-PD-1 response. The expression of BHLHE40 by tumor-specific cytolytic T cells may increase the recruitment of macrophages. In turn, these macrophages, together with DCs, may increase the recruitment of tumor-specific and stem-like TCF+ T cells through antigen presentation by MHC and CXCL9 production. Finally, the expression of CCL5 by tumor-specific and stem-like TCF+ T cells and NK may increase the recruitment of additional T cells.

### Immunosuppressive tumor and myeloid cells, along with the expression of multiple checkpoints, characterize tumor resistance

We next aimed to identify immunosuppressive pathways exhibiting resistance to anti-PD-1 therapy through single-cell RNAseq analysis of tumor cells. MSH2 KO and WT tumor cells clustered distinctly, with only a limited number of clusters shared between these two tumor types (**Extended Figure 10**). MSH2 KO tumors generated a more diverse array of clusters compared to WT tumors (12 vs 4, respectively), characterized by a higher expression of MHC and B2M (**Extended Figure 10**). Interestingly, anti-PD-1 therapy led to a reduction in the cell count within MSH2 KO cluster 6 and WT cluster 7, suggesting that these clusters may be targeted by immune cells (**Extended Figure 10**). Post anti-PD-1 treatment, a surge of cells emerged in MSH2 KO cluster number 8 and WT cluster 1, implying the resistance of these clusters to the therapy (**Extended Figure 10**). Within these resistant clusters, an upregulation of genes such as IFITM2, IFITM3, TMEM176, regulating inflammasome and epithelial-mesenchymal transition ^31–33^, and other resistance-associated genes were detected when juxtaposed with clusters effectively targeted by anti-PD-1 (**Extended Figure 10**). Gene enrichment analysis in these tumor scRNAseq clusters indicated that resistance involved pathways similar to the ones that negatively regulate viral entry into host cells (**Extended Figure 11**). In addition, VEGFB exhibited higher expression in MSH2 KO tumors when contrasted with WT tumors (**Extended Figure 10 B**), a phenomenon corroborated by our findings through luminex analysis on spheroids (**Extended Figure 12 C**). Overall, tumor cells may escape from immune surveillance by upregulating genes associated with resistance in multiple cancers such as IFITM and TMEM176.

In addition to utilizing intrinsic immunosuppressive tactics, tumors can alter the immune milieu to drive a pro-tumorigenic inflammation. To explore whether the immune escape of MSH2 KO tumors might be linked to immune checkpoint expression, T cell exhaustion, or myeloid immunosuppressive programs, we quantified PD-1, TIM3, TIGIT, CTLA4 and LAG3 immune checkpoints, as well as their corresponding ligand expression in tumor-infiltrating immune cells using flow cytometry and scRNAseq. Through scRNAseq, we identified high levels of PD-1, TIM3, TIGIT, CTLA4, and LAG3 expression in CD8+ TILs (**Figure 3 A**). Post anti-PD-1 therapy, the myeloid-infiltrating cells exhibited elevated expression of checkpoint ligands, particularly notable for LAG3 ligands, and molecules such as TREM2, CSF1R, and IL1B, a monocyte cytokine previously associated with resistance in MMRd CRC tumors ^17^. Additionally, an increase in infiltrating monocytes and macrophages was observed (**Figure 3 A, Extended Figure 3**). PD-L1 and TIM3 ligands (LGALS9, PTDSS1, CEACAM1, and HMGB1) were identified on macrophages, DCs, and tumor cells. TIGIT ligands (PVR, NECTIN2, NECTIN3) were predominantly expressed on tumor cells, while LAG3 ligands (H2-AB1, H2-AA, H2-EB1) were mainly expressed on macrophages and DCs. Notably, TREM2 and CSF1R showed prevalent expression in macrophages, IFITM2, and 3 in monocytes, and IL1B in neutrophils and monocytes. Hence, overexpression of immunosuppressive molecules by myeloid cells and checkpoint expression by T cells could potentially trigger ICB resistance to anti-PD-1 (**Figure 3 A**). Using spectral flow cytometry, we verified an increase in CD8+ T cells expressing PD-1, TIM3, and TIGIT in MSH2 KO tumors when compared to WT (**Figure 3 B**). MSH2 KO tumors displayed a higher presence of exhausted PD-1+ TCF-TIM3+ T cells (TEX) but not their PD-1+ TCF+ T cell progenitors (TPEX) (**Figure 3 B, Extended Figure 2**). Notably, the elevated PD-1 expression on CD8+ T cells might elucidate the efficacy of anti-PD-1 therapy in MMRd patients. After 28 days, we noted an escalation in CD8+ T cells expressing TIM3 and TIGIT in MSH2 KO tumors treated with anti-PD-1 compared to untreated MSH2 KO and WT tumors (**Figure 3 B**). Furthermore, an increase in neutrophil, macrophage, and cDC2 infiltration was observed in MSH2 KO tumors following anti-PD-1 treatment (**Figure 1 C**). In MSH2 KO tumors treated with anti-PD-1, the spleen and lymph nodes also exhibited an increased presence of CD8+ T cells expressing LAG3, TIGIT, TIM3, and myeloid cells (**Figure 3 C, Extended Figure 6 D**). Overall, the immune resistance observed in MMRd tumors after anti-PD-1 treatment might be induced by the expression of other immune checkpoints like TIM3, LAG3, CTLA4, TIGIT, as well as the expression of TREM2, CSF1R, IFITM, and IL1B by myeloid cells. These resistance programs appear to involve the increased production of these cells by the spleen and their recruitment from the lymph nodes.

**Figure 3.**
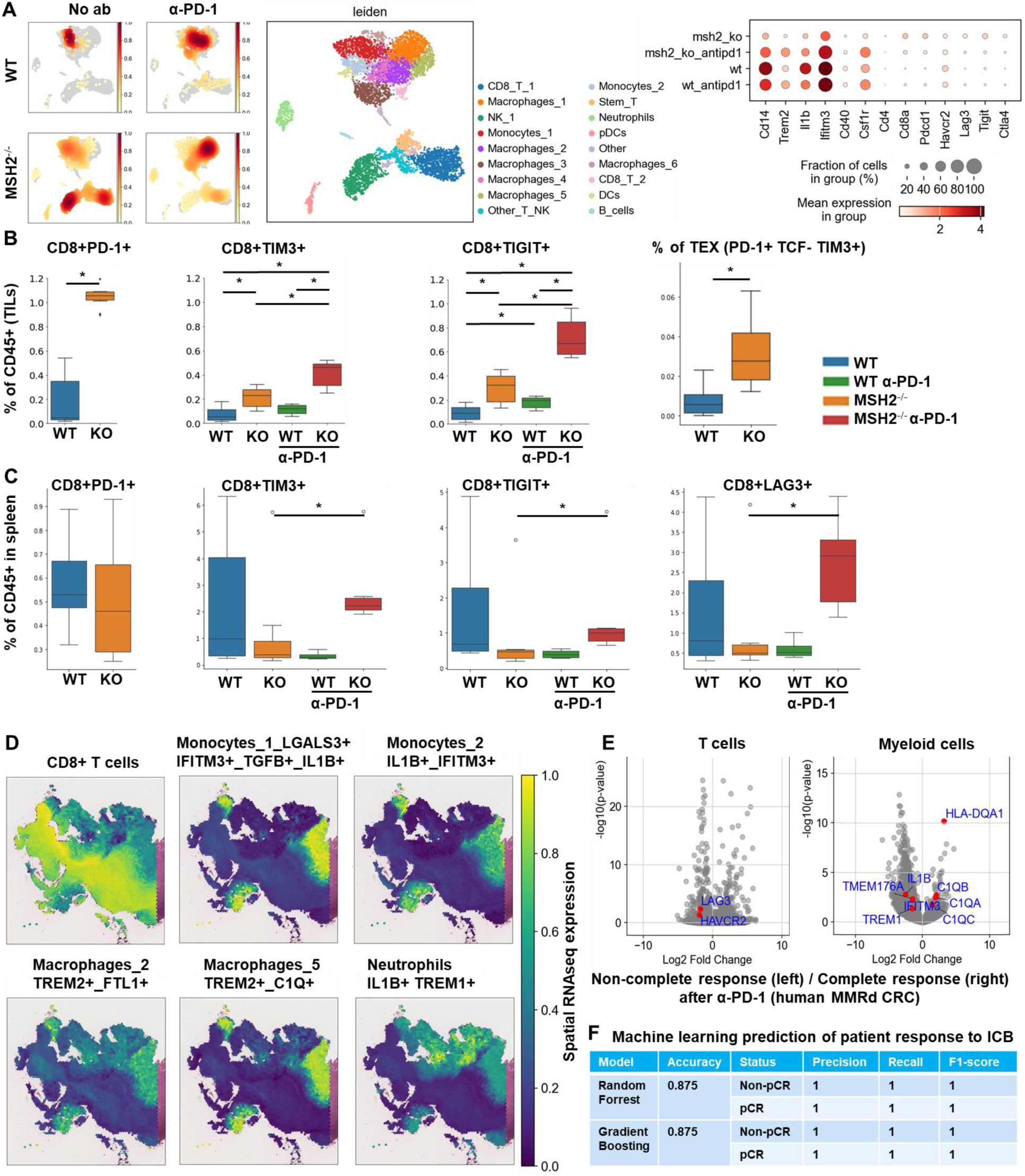
Immunosuppressive myeloid cells, along with the expression of multiple checkpoints, characterize tumor resistance. **A** ScRNAseq UMAP of immune cells was generated, following the exclusion of tumor cells, in both CT26 WT and MSH2 KO tumors, with and without anti-PD-1 therapy. The distribution of cells and immune genes in each tumor cluster is depicted. **B/C** The quantification of differential immune checkpoint and lymphocyte infiltration expression in tumors (**B**) and spleens (**C**), contingent on the MSI status and ICB, is shown through flow cytometry after 28 days. N=6 replicates, Mann-Whitney U Test, p value < 0.05. Data are represented as mean ± SEM. **D** Spatial localization of each single-cell immune cluster in MSH2 KO tumors with anti-PD-1 therapy. The CD8 panel is the same as in Figure 2 E. **E** A reanalysis of previously published human MMRd CRC patient data ^17^ has been conducted, focusing on patients with complete or incomplete responses to anti-PD-1 therapy alone, and scrutinizing the underlying T cell and macrophage profiles. Gene expression within each immune cluster. **F** RandomForestClassifier and GradientBoosting algorithms trained on a scRNAseq dataset ^17^, n=10 MMRd CRC patients. Accuracy = correct predictions / total number of predictions. Precision = correct predictions of a class / all positive true positive and false positive predictions. Recall (Sensitivity) = correct true positive predictions of a class / actual instances of the class. F1-score = harmonic mean of precision and recall.

Our next objective was to determine if the resistance to anti-PD-1 involving multiple checkpoints on TILs depends on interactions with the TME, or is mediated by a direct effect of PD-1 blockade on naive immune cells recruited to the tumor. To achieve this, we constructed tumor models with no TME, which are CT26 and B16F10 spheroids comprising MSH2 KO or WT cells, and co-cultured them with naïve splenic cells along with anti-PD-1 treatment. Utilizing 3D bi-photon imaging, we observed the infiltration of naïve splenic cells into both WT and MSH2 KO CT26 spheroids post anti-PD-1 administration (**Extended Figure 12 A**). These findings were corroborated by our flow cytometry analysis (**Extended Figure 12 B/D**). Upon the supplementation of naïve splenic cells with anti-PD-1 or anti-CTLA4, we noticed an escalation in CD8+ T cells expressing CD25, CD69, Ki67, TIM3, and TCF, accompanied by a decline in Tregs (**Extended Figure 12 B**). The upregulation of other checkpoints on CD8+ T cells following ICB, such as LAG3 and TIM3 was also observed in the B16F10 model (**Extended Figure 12 D**). Consequently, the immune resistance of MMRd tumors following ICB appears to involve the early upregulation of other immune checkpoints, such as TIM3 and LAG3, due to PD-1 blockade direct effect on immune cells, and is not always dependent on the TME.

To investigate if there were immunosuppressive mechanisms involving interactions between T cells and myeloid cells in the context of the TME, we applied scRNAseq clustering to spatial samples. We noted the co-localization of NK cells and monocytes expressing elevated IFITM, LGALS3, TGFB, and IL1B, alongside macrophages in clusters 2, 5, and 6 showcasing heightened TREM2 and IFITM levels. These clusters were found within the tumor core, notably within zones marked by the exclusion of CD8 T cells (**Figure 3 A/D, Extended Figure 3/9**). Neutrophils with elevated IL1B and TREM1 levels were also identified in areas marked by CD8+ T cell exclusion after anti-PD-1 therapy **(Figure 3 A/D, Extended Figure 3/9**). Immunosuppressive macrophages clusters 2, 5 and 6, monocytes and neutrophils express high levels of VEGF and IL1RN, which are cytokines that promote Tregs and limit inflammation through competition with IL1, respectively (**Extended Figure 3 D**). Collectively, resistance in WT tumors appears to be orchestrated by the infiltration of monocytes and the expression of immunosuppressive molecules like IFITM, LGALS3, TGFB, IL1B, VEGF and IL1RN. Conversely, resistance in MSH2 KO tumors could potentially stem from immunosuppressive macrophages expressing TREM2, VEGF and IL1RN expression, TREM1+ neutrophils producing IL1B, and the presence of multiple T cell checkpoints, including TIM3 and LAG3.

Finally, we undertook a reanalysis of previously published human MMRd CRC patient data ^17^, with a particular emphasis on patients who had a complete responses or no response to anti-PD-1 therapy, while scrutinizing the underlying profiles of myeloid and T cells (**Figure 3 E, Extended Figure 5**). Notably, human MMRd CRC patients who did not respond to anti-PD-1 treatment demonstrated an increased presence of myeloid cells expressing elevated levels of IL1B, TREM1, IL1RN, IDO1, IL6, TMEM176 and IFITM (**Figure 3 E, Extended Figure 5**), akin to our observations in the murine model. Furthermore, non-responders also exhibited a higher population of CD4+ and CD8+ T cells displaying heightened levels of TIM3 and LAG3 (**Figure 3 E, Extended Figure 5**), in parallel to our observations in mice. We hypothesized that we could use these scRNAseq data to predict which patients will resist or respond to ICB. By applying RandomForrest and GradientBoosting machine learning algorithms on these scRNAseq datasets, divided into training and validation datasets, we correctly predicted the response to ICB for 87.5% of the patients (**Figure 3 F**). By reanalyzing mutational data from 2 other human CRC cohorts, we observed that most of the non-responders are characterized by a MSS status ^13,34^. By applying machine learning algorithms on these datasets, we predict ICB response for these patients (**Extended Figure 13**). Thus, machine learning can successfully capture the complexity of individual patient profile to correctly predict the response. Overall, these results imply that targeting myeloid subsets and employing multiple checkpoint blockade strategies might hold potential benefits.

### Combination of TREM2, LAG3, CTLA4 and PD1 blockade eliminates both MMRd and MMRp tumors

Based upon our observations, we next investigated the impact of targeting T cell checkpoints on tumor growth by using anti-TIM3, anti-TIGIT, anti-LAG3, or anti-CTLA4, alone or in combination with anti-PD-1. We injected CT26, 4T1 and B16F10 MMRd or MMRp cell lines in mice and used anti-TIM3/TIGIT/LAG3/CTLA4 alone or in combination with anti-PD-1, 14 days following tumor inoculation. The in vivo tumor volume was assessed and compared, taking into consideration both immune checkpoint blockade and MSH2 status (**Figure 4 A/C/F/H, Extended Figure 14, Extended Figure 15 A/C/F/H**). Importantly, upon the administration of anti-TIM3/TIGIT/LAG3, or anti-PD-1, we observed suppression in the growth of MSH2 KO CT26 tumors after 28 days (**Figure 4 C, Extended Figure 15 C**).

**Figure 4.**
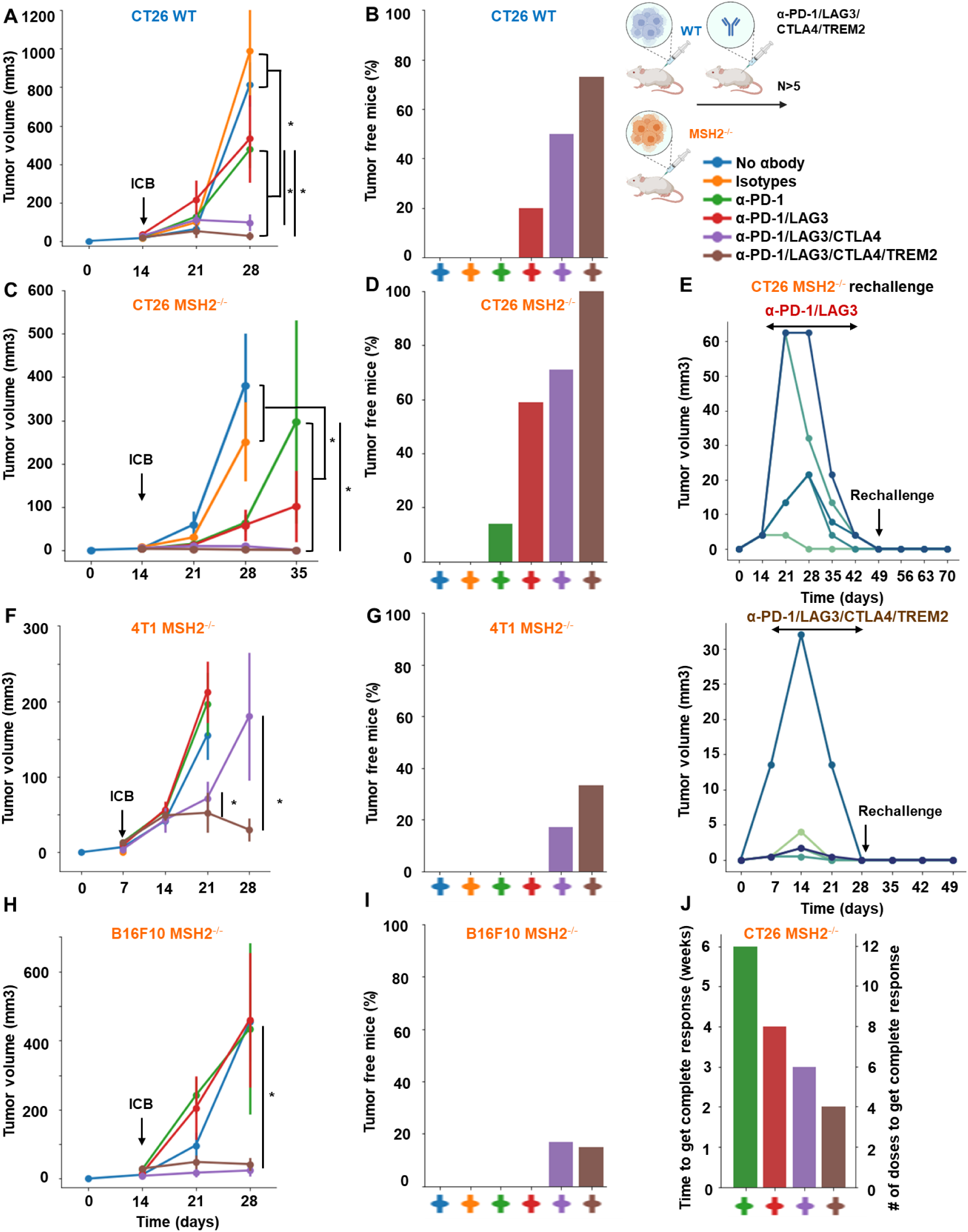
Combination of TREM2, LAG3, CTLA4 and PD1 blockade eliminates both MMRd and MMRp tumors and elicit robust immune memory. **A/C/F/H** Tumor volume (mean) in vivo over time of CT26, B16F10 and 4T1 WT and MSH2 KO cells (200,000 cells/tumor) in the absence or presence of anti-PD-1, anti-LAG3, anti-TREM2 or anti CTLA4 (100 µg twice a week after 14 days) therapy (in mm3). N= 5 to 20 per arm. Mann-Whitney U Test, p value<0.05. Data are represented as mean ± SEM. **B/D/G/I** Complete response of CT26, B16F10 and 4T1 WT and MSH2 KO tumors after ICB combinations (%). **E** After CT26 MSH2 KO tumor elimination following anti-PD-1 + anti-LAG3 +/- anti-TREM2 therapy, treatment was stopped and a second tumor was inoculated on the opposite flank and its growth was measured for 3 weeks. N= 5. Each curve is for an individual mouse. **J** Time and number of antibody doses needed to observe complete response and CT26 MSH2 KO tumor elimination.

This effect was further amplified with the utilization of anti-TIM3/TIGIT/LAG3/CTLA4 in combination with anti-PD-1 after 35 days, with the most favorable outcomes seen for the anti-PD1/LAG3/CTLA4 combination. These combinations led to an increase in complete responses and even tumor elimination (**Figure 4 C/D**, **Table 1, Extended Figure 15 C/D**). WT tumors exhibited responsiveness to the anti-PD-1/LAG3, the anti-PD-1/TIGIT and anti-PD1/LAG3/CTLA4 combinations (**Figure 4 A/B, Extended Figure 15 A/B**). In the context of MMRd CRC tumors, we observed complete responses of 25%, 12%, 59% and 71% for anti-PD-1/TIM3, anti-PD-1/TIGIT, anti-PD-1/LAG3 and anti-PD1/LAG3/CTLA4 respectively (**Figure 4 D, Extended Figure 15 D**). On the other hand, for WT CRC tumors, complete responses were observed in 0%, 10%, 20% and 50% cases for anti-PD-1/TIM3, anti-PD-1/TIGIT, anti-PD-1/LAG3 and anti-PD1/LAG3/CTLA4 respectively (**Figure 4 B, Extended Figure 15 B**). Incorporating the 4T1 and B16F10 models, we also noted a restriction in MMRd tumor growth upon administration of anti-PD-1/CTLA4/LAG3 (**Figure 4 F/H, Extended Figure 15 F/H**). Interestingly, to observe a complete response in the 4T1 and B16F10 models, it was essential to include anti-CTLA4 alongside the combinations identified for the CT26 model (**Figure 4 G/I, Extended Figure 15 G/I**).

We also investigated the impact of targeting myeloid cell checkpoints together with T cell checkpoints on tumor growth by using anti-IL1B, anti-TREM2 or anti-IFITM, alone or in combination with anti-PD-1/CTLA4/LAG3. Targeting TREM2 or IFITM limited MMRd tumor growth (**Figure 4 C, Extended Figure 15 C**), but blocking IL1B was not beneficial. Blocking TREM2 in addition to T cell checkpoints further limited tumor growth in MMRd and MMRp models (**Figure 4 A/B/C/D/F/G, Extended Figure 15, Table 1**). In the context of MMRd CRC tumors, we observed complete responses of 20%, 50% and 10% for anti-PD1/TREM2, anti-PD1/LAG3/TREM2 and anti-PD1/IFITM respectively (**Figure 4 D, Extended Figure 15 D**). Notably, the most favorable outcomes were seen for the anti-PD1/LAG3/CTLA4/TREM2 combination, with complete responses of 100% for MMRd CRC and 73% for MMRp CRC (**Figure 4 B/D, Extended Figure 15)**. This combination was also efficient in B16F10 and 4T1 models (**Figure 4 F/G/H/I, Extended Figure 15)**. The number of antibody doses and time needed to achieve complete response and tumor elimination was reduced when using multiple ICB and anti-TREM2 compared to anti-PD-1 alone (**Figure 4 J**), limiting mice exposure to potential chronic adverse events. Overall, the combination of TREM2, LAG3, CTLA4 and PD1 blockade is the most efficient and eliminates both MMRd and MMRp tumors.

Next, we investigated if tumor elimination following multiple checkpoint blockade successfully elicited a memory response and could prevent tumor recurrence. We observed that all mice were effectively protected against a second tumor inoculation subsequent to the elimination of the initial tumor through multiple ICB, suggesting the establishment of potent anti-tumor immune memory mediated by the combination of ICB treatments (**Figure 4 E**). Furthermore, we noted that the proportion of effector memory CD8+ and CD4+ T cells in the spleens and lymph nodes of mice that successfully eradicated an MSH2 KO tumor following anti-PD-1 and anti-LAG3 therapy was higher compared to mice unable to eliminate the tumor (**Extended Figure 15 J**). In conclusion, these findings underscore that targeting multiple immune checkpoints and TREM2 macrophages in MMRd and MMRp tumors not only further curbs tumor growth but can also culminate in complete tumor eradication. The utilization of such ICB and myeloid targeting combinations holds potential for mitigating immune resistance in both MMRp and MMRd tumors.

### TCF+ T cells, CD4+ and CD8+ T cells, MHC+ macrophages and neutrophils orchestrate the response to targeted checkpoint/myeloid combinations

To evaluate if the impact of targeted ICB on MSH2 KO tumor growth limitation was due to immune infiltration, we examined the immune subsets that differentially infiltrate MSH2 KO and WT tumors in mice after anti-TIM3/TIGIT/LAG3, with or without anti-PD-1 therapy (**Extended Figure Figure 16 A/B, Extended Figure 17 A/E**). We compared immune infiltration in tumors based on tumor weight and ICB status (**Extended Figure 16 A**). Notably, we observed weight reduction in MSH2 KO tumors after multiple ICB interventions. Crucially, there was no discernible difference in overall immune infiltration correlated with tumor weight. We detected increased infiltration of CD8+ T cells, particularly those expressing TCF, KI67, PD1, TIM3, LAG3, and TIGIT, alongside CD4+ T cells, mast cells, macrophages, neutrophils in smaller tumors, coupled with increased MHC expression (**Extended Figure 16 A**). Additionally, anti-LAG3 increased MHCII expression on tumors, reduced myeloid infiltration, and enhanced T cell infiltration. Anti-TIM3 decreased myeloid infiltration and increased CD4+ TCF+ T cell infiltration. Anti-TIGIT increased both CD4+ TCF+ T cell and CD8+ TCF+ T cell infiltration (**Extended Figure 17 A**). By scRNAseq and spatial RNAseq, we observed that following anti-PD-1 + anti-CTLA4 + anti-LAG3 + anti-TREM2, MSH2 KO tumors were more infiltrated by CD4 T cells, MHC+ macrophages, neutrophils, TCF+ or ICOS+ T cells and less SPP1+ macrophages, previously described as immunosuppressive ^35^, monocytes and exhausted T cells compared with untreated or anti-PD-1 treated mice (**Figure 5 A/B, Extended Figure 9**). Importantly, when adding anti-TREM2 to PD1/LAG3 blockade, mice with a decrease in tumor volume had tumors less infiltrated by immunosuppressive macrophages, while still enriched in TCF+ PD1+ T cells and DCs compared to non responders (**Figure 5 C**). Compared to CT26 MSH2 KO tumors, 4T1 MSH2 KO tumors contained more monocytes and neutrophils but less T cells, especially T cells expressing TCF and PD1, and that may explain their stronger resistance to ICB (**Extended Figure 18**). By measuring T cell phenotype and MMRd spheroid killing following coculture with MonoMacs from tumor or LN of mice bearing CT26 MSH2 KO tumors treated with anti-PD-1 or anti-PD-1/LAG3/CTLA4/TREM2, we observed an increase in T cell activation and tumor spheroid killing following coculture with MonoMacs from mice treated with anti-PD-1/LAG3/CTLA4/TREM2 (**Extended Figure 19**). Overall, response to multiple checkpoint and myeloid blockades is driven by both CD4 and CD8 T cells, including T cells expressing TCF, ICOS, PD-1 and KI67, as well as neutrophils and MHC+ macrophages, and MHC expression by tumor cells.

**Figure 5.**
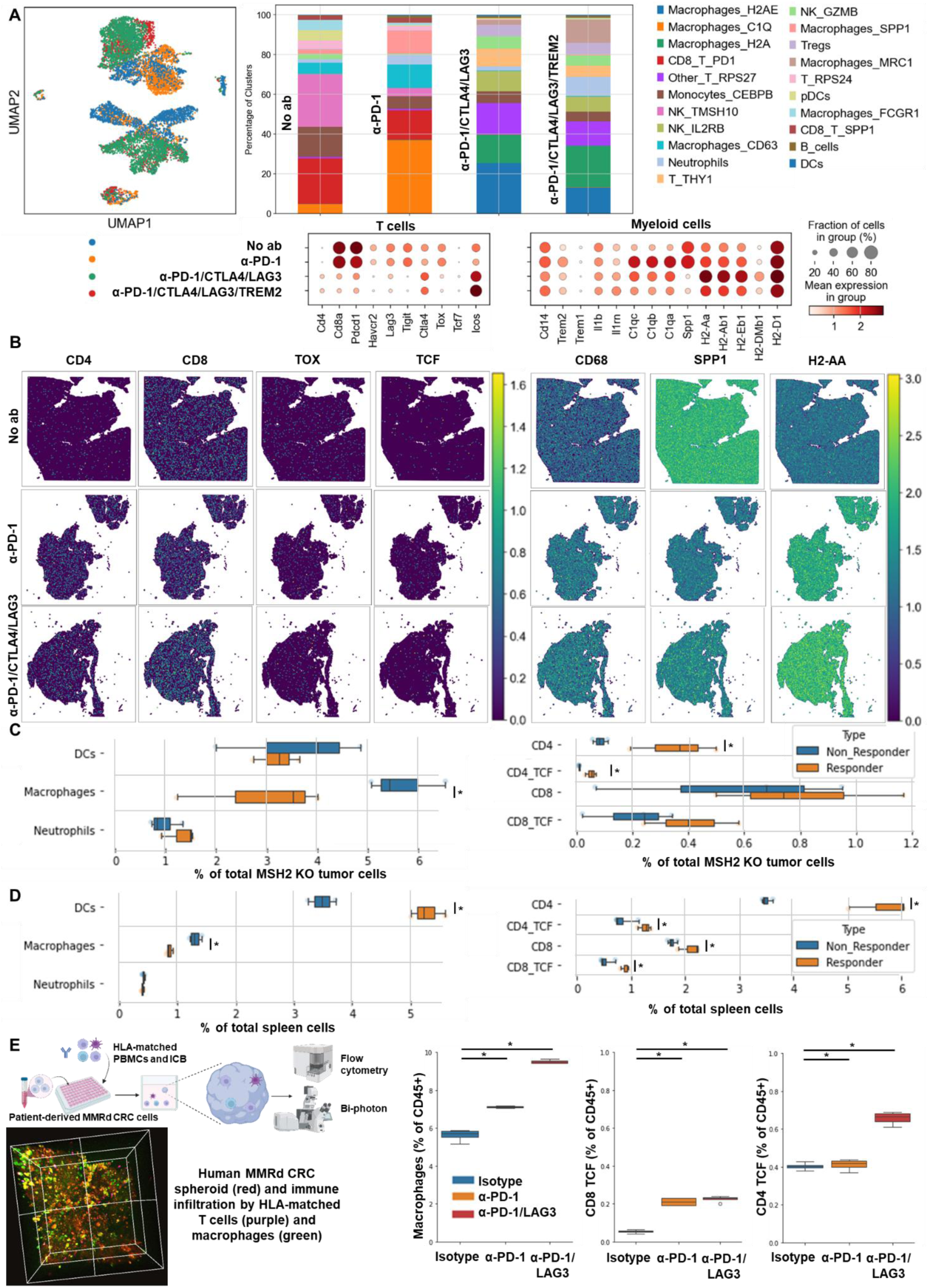
TCF+ T cells, CD4+ and CD8+ T cells, neutrophils and MHC+ macrophages orchestrate the response to targeted checkpoint/myeloid combinations. CT26 tumors deleted for MSH2 are growing for 28 days in BALB/C mice. **A** ScRNAseq UMAP of immune cells in CT26 MSH2 KO tumors with or without anti-PD-1/CTLA4/LAG3/TREM2 therapy. UMAP of immune clusters subsequent to Leiden clustering by scanpy. Percentage of cells in each immune cluster and gene expression within each immune cluster. **B** Spatial RNAseq localization of each single-cell immune cluster in MSH2 KO tumors following ICB. **C** Quantification of differential immune checkpoint and lymphocyte infiltration expression according to response to PD1/LAG3/TREM2 blockade. N=8 replicates, t-Test, p value<0.05 Data are represented as mean ± SEM. Mice with a decrease in tumor volume following therapy are considered as responders. **D** Immune composition in spleens and lymph nodes was also measured (flow cytometry). N= 8. Mann-Whitney U Test, p value<0.05. Data are represented as mean ± SEM. **E** 10,000 MMRd CRC patient-derived cells are growing for 3 days as spheroids and 100,000 PBMC from 3 donors are activated with IL-15. Then, spheroids and PBMC are incubated for 3 days. PBMC were incubated before with anti-PD-1, anti-TIM3 and anti-LAG3 (10 µg/mL). Immune cells are previously labeled by cell tracker (red). Spheroid cell nuclei are labeled with Nunc blue (blue). 3D Bi-photon imaging and flow cytometry staining of spheroids after dissociation. Quantification of differential immune checkpoint expression and lymphocyte infiltration according to ICB. N=4. Mann-Whitney U Test, p value<0.05 Data are represented as mean ± SEM.

We explored whether this increase in immune infiltration of MSH2 KO tumors following multiple ICB/myeloid targeting therapy might be attributable to an increased immune recruitment from lymph nodes and spleen. To investigate this, we assessed immune distribution through flow cytometry in the spleen and lymph nodes 28 days post-tumor injection and multiple ICB/myeloid targeting therapy. We compared these outcomes in mice that responded (tumor rejection) and those that did not (**Extended Figure 16 B/C, Extended Figure 17 C/D/F**). Responders to multiple ICB/myeloid targeting had increased counts of CD4+ T cells, CD8+ T cells, TCF+ T cells, CD8+ T cells expressing KI67, PD-1, DCs, along with diminished mast cells, basophils, and polymorphonuclear MDSCs in both the spleen and lymph nodes (**Figure 5 D**, **Extended Figure 16 B/C, Extended Figure 17 C/D/F**). Blocking IL1B was not beneficial and was associated with a diminution in TCF+ T cells and an increase in Tregs in the spleen (**Extended Figure 16 F**). Thus, the increased infiltration of T cells and especially T cells expressing TCF, PD1 and KI67 into MSH2 KO tumors following successful multiple checkpoint blockade might be attributed to escalated recruitment from both spleen and lymph nodes.

To explore the effectiveness of ICB combinations in a human context, we conducted experiments using patient-derived MMRd CRC spheroids with immune infiltration and stimulation using anti-PD-1, anti-TIM3, and anti-LAG3. We observed that ICB led to an increased infiltration of TCF+ T cells, along with MHC+ macrophages and neutrophils (**Figure 5 E, Extended Figure 19**). Notably, the infiltration of these subsets was further amplified when anti-PD-1 was combined with either anti-TIM3 or anti-LAG3. These findings align with the results obtained from the mouse model (**Figure 5 A/B/C**), suggesting that ICB combinations could also prove effective in human contexts and involve TCF+ T cells, both CD4 and CD8 T cells, neutrophils and MHC+ macrophages.

## Discussion

ICB therapy demonstrates a high response rate and durable clinical benefit in patients with MMRd tumors. Our study revealed that the immune response of MMRd CRC tumors is mediated by myeloid and lymphocyte coordination, including T cells, DCs, NK cells, and MHC+ C1Q+ macrophages. Additionally, we found that this broad immune response is enhanced following anti-PD-1 therapy in both mouse and human settings. T cells may recognize highly immunogenic shared frameshift-derived neoantigens in MMRd tumors ^13^. The observed overexpression of PD-1 in MMRd tumors might explain why they are more responsive to anti-PD-1 therapy ^6–9^. Previous research demonstrated the significance of TCF+ T cells in the response to anti-PD-1 in a melanoma model ^21^, and our results indicate a similar mechanism in MMRd CRC tumors. Intriguingly, we demonstrated that the response also involves MHC+ C1Q+ CXCL9+ macrophages and DCs that co-localize between T cell exclusion zones and T cell-enriched zones. Notably, this C1Q signature was linked to a poor prognosis in clear-cell renal cell carcinoma ^22^, but CXCL9 attracts CD8+ cytotoxic T cells and is associated with better outcomes ^23^. These results are in line with recent data from human cohorts that suggest that responsive hypermutated CRCs were enriched in cytotoxic and proliferating PD1+ CD8 T cells interacting with PDL1+ antigen-presenting macrophages ^34^. The remodeling of the infiltrating monocytes and macrophages was observed in other models following ICB ^36^. Thus, the collaboration among DCs, MHC+ C1Q+ macrophages, and CD8+ BHLHE40+ TCF+ T cells plays a crucial role in the anti-PD-1 response in MMRd tumors. This response is facilitated by an increased production of these cells by the spleen and their recruitment from the lymph nodes.

40-70% of patients with advanced MMRd tumors do not respond to anti-PD-1 therapy ^1,11,12^. We have demonstrated that MMRd tumor resistance involves immunosuppressive tumor and myeloid cells, as well as the expression of multiple checkpoints. Our observations indicate that MMRd tumors possess a higher transcriptomic diversity compared to MMRp tumors, and anti-PD-1 therapy does not effectively target all these tumor clusters. Those clusters that resist anti-PD-1 treatment upregulate IFITM, TMEM176, controlling inflammasome and epithelial-mesenchymal transition, and other genes associated with resistance in multiple cancers ^31–33,37^. Furthermore, we have observed that tumors resistant to anti-PD-1 treatment exhibit higher infiltration of TREM2+ macrophages, monocytes, neutrophils, and monocytes expressing IL1B. These subsets colocalize in T cell exclusion zones. Previously, high expression of IL1B by monocytes has been associated with resistance in MMRd CRC tumors ^17^, as well as TREM2 in multiple models ^24–27^. Other mechanisms of resistance may involve cancer-associated fibroblasts, Tregs, and neutrophils ^16,17,38^. Importantly, our data demonstrate that the immune resistance of MMRd tumors can also be attributed to the expression of multiple immune checkpoints (PD-1, TIM3, LAG3, CTLA4, TIGIT). This finding aligns with the fact that MMRd polyps are characterized by CTLA-4, LAG3, and PD-L1 expression ^14–16^. The resistance of MMRp tumors is mediated by monocytes that highly express IFITM, TMEM176B, LGALS3, TGFB, and IL1B. In MMRd tumors, resistance following anti-PD-1 treatment is driven by the expression of TIM3, LAG3, CTLA4, and TIGIT by TILs, as well as tumor clonal selection and the expression of TREM2, CSF1R, IFITM, TMEM176B and IL1B by myeloid cells. These resistance programs involve an increased production of these immune cells by the spleen and presence in the lymph nodes.

Our data showed that checkpoint receptor and myeloid cell targeting combinations overcome MMRp and MMRd tumor resistance. Following anti-TIM3, anti-TIGIT, anti-LAG3, anti-TREM2 or anti-PD-1 treatment, we observed a limitation in MSH2 KO CRC tumor growth. This effect was amplified when using combinations of anti-PD1 with anti-TIM3/TIGIT/ LAG3/TREM2/IFITM/CTLA4. The most robust responses were observed for the anti-PD1/LAG3/CTLA4/TREM2 combination in MMRd tumors and MMRp tumors. Notably, this combination significantly increased the occurrence of complete tumor elimination, reaching up to 100% for MMRd CRC tumors and up to 73% for MMRp CRC tumors. Adding anti-CTLA4 to the combinations identified in the CRC model was necessary to achieve a complete response in breast and melanoma MMRd cancer models, which is consistent with other studies ^38^. We observed that all mice that successfully eliminated a first tumor through targeted checkpoint therapy were effectively protected against a second tumor inoculation, indicating the development of effective anti-tumor immune memory, and may limit the risk of neoplasia ^39^. Thus, tailored immunotherapies based on the patient’s tumor profile hold potential benefits.

By measuring immune infiltration in mouse tumors, spleens, lymph nodes, and human MMRd CRC patient-derived spheroids, we observed that the response to targeted checkpoint blockade involves MHC+ macrophages, both CD4 and CD8 T cells, neutrophils, TCF+ T cells, and memory T cells. This response also correlates with a reduction of T cell exhaustion and infiltration by SPP1+ TREM2+ macrophages. Consistently with our results, T cells expressing TCF have been shown to be a stem-like population that is central to the maintenance of long-term antiviral immunity and responsiveness to immunotherapy in a melanoma model ^21,40^. Our results showed that this stem T cell subset is also key in the response to multiple checkpoint blockade, especially in tumors with high indel and snv loads, together with MHC+ macrophages and neutrophils.

Our findings have implications for cancer immunotherapy. Our results show that strategically targeting TIM3, LAG3, TIGIT, CTLA4, IFITM, TREM2 and PD-1 in MMRd and also MMRp CRC tumors not only further limits tumor growth but can also significantly increase the complete elimination of tumors. Using anti-LAG3 or anti CTLA4 with anti-PD-1 is promising for MMRd CRC patients ^41,42^ and other cancers such as melanoma ^43,44^. We showed that targeting these 3 checkpoints together and targeting myeloid cells based on the patient’s tumor profile could aid in mitigating immune resistance in both MMRp and MMRd tumors. These combinations could be employed alongside other strategies targeting myeloid subsets, including monocytes, neutrophils and immunosuppressive macrophages ^24,26,45,46^.

While we have provided comprehensive analyses using orthogonal approaches in both human and mouse settings, it is likely that TCF+ T cells, MHC+ macrophages, and neutrophils are not the sole subsets driving targeted ICB responses in MMRd CRC tumors. Complementary approaches could also focus on γδ T cells, especially in MMRd cancers with MHCI defects ^47^, DCs ^29,48^, PD-1 and PI3K-γ expressing myeloid cells ^49,50^, the HLA-E and HLA-G molecules ^20,35,51–56^, WRN inhibitors ^57^ or TREM1 myeloid cells ^58^. In conclusion, our results strongly indicate that TCF+ T cells and MHC+ macrophages function as key effector cells in targeted ICB/myeloid therapy for MMRd CRC.

## Methods

### Key Resources Table

**Table 1.** List of antibodies.

| **<u>Table 1) List of antibodies.</u>** |  |  |
| --- | --- | --- |
| <u>Target</u> | <u>Fluorochrome</u> | <u>Supplier</u> |
| anti-CD3 Rat | Alexa Fluor 488 | BioLegend |
| anti-mouse CD4 [GK1.5] | Alexa Fluor 532 | Thermo Scientific |
| anti-mouse F4/80 [BM8] | Alexa Fluor 532 | eBioscience |
| anti-mouse H-2Dd [34-2-12] | PE (Phycoerythrin) | BioLegend |
| anti-CD370 (CLEC9A) Rat [7H11] | PE (Phycoerythrin) | BioLegend |
| anti-CD25 Rat [PC61] | Pe/dazzle™ 594 | BioLegend |
| anti-mouse CD86 [GL-1] | Pe/dazzle™ 594 | BioLegend |
| anti-CD11b Rat [M1/70] | PerCP | BioLegend |
| anti-TIGIT [GIGD7] | PerCP-eFluor 710 | Thermo Scientific |
| anti-CD274 (PD-L1) Rat [10F.9G2] | PE (Phycoerythrin)/Cy7® | BioLegend |
| anti-mouse I-A/I-E [M5/114.15.2] | APC | BioLegend |
| anti-mouse CD45 | Spark NIR™ 685 | BioLegend |
| anti-mouse TCF-7/TCF-1 [S33-966] | R718 | BD |
| anti-Ly-6C Rat [HK1.4] | Alexa Fluor® 700 | BioLegend |
| anti-mouse FcεRIα | APC/Fire™ 750 | BioLegend |
| anti-mouse CD69 [H1.2F3] | APC/Cy7 | BioLegend |
| anti-mouse CD279 (PD-1) [J43] | Super Bright 436 | Thermo Scientific |
| anti-mouse CD206 (MMR) [MR6F3] | eFluor 450 | Thermo Scientific |
| anti-FOXP3 Rat [MF-14] | Pacific Blue | BioLegend |
| anti-CD335 Rat [29A1.4] | Brilliant Violet® 510 | BioLegend |
| anti-CD103 Armenian Hamster [2E7] | Brilliant Violet® 510 | BioLegend |
| anti-CD19 Rat [6D5] | Brilliant Violet® 570 | BioLegend |
| anti-CD11c Armenian Hamster [N418] | Brilliant Violet® 570 | BioLegend |
| anti-rat CD8a [53-6.7] | Brilliant Violet® 605 | BioLegend |
| anti-mouse/human Ki-67 | Brilliant Violet 650™ | BioLegend |
| anti-mouse CD117 (c-kit) | Brilliant Violet 650™ | BioLegend |
| anti-mouse CD366 (Tim-3) | Brilliant Violet 711™ | BioLegend |
| anti-rat Ly-6G/Ly-6C (Gr-1) [ RB6-8C5] | Brilliant Violet® 711 | BioLegend |
| anti-mouse Ly-6G [1A8] | Brilliant Violet 785™ | BioLegend |
| anti-mouse CD223 (LAG-3) [C9B7W] | Brilliant Violet 785™ | BioLegend |
| anti-human CD16 | NovaFluor Blue 510 | Biolegend |
| anti-human CD11c | FITC | Biolegend |
| anti-human FOXP3 | AlexaFluor 532 | Biolegend |
| anti-human CD19 | NovaFluor Blue 585 | Biolegend |
| anti-human CD4 | NovaFluor Blue 610-70S | Biolegend |
| anti-human CD45 | PerCP Cy5.5 | Biolegend |
| anti-human HLA-ABC | PercCP-eFluor 710 | Biolegend |
| anti-human TIGIT | BV421 | Biolegend |
| anti-human ILT2 | SB436 | Biolegend |
| anti-human CD3 | Pacific Blue | Biolegend |
| anti-human CD11b | BV480 | Biolegend |
| anti-human CD25 | BV510 | Biolegend |
| anti-human CD56 | BV570 | Biolegend |
| anti-human FCER1A | BV605 | Biolegend |
| anti-human CD86 | BV650 | Biolegend |
| anti-human CD68 | BV711 | Biolegend |
| anti-human CD15 | BV750 | Biolegend |
| anti-human LAG3 | BV786 | Biolegend |
| anti-human TCF1 | AlexaFluor 647 | Biolegend |
| anti-human HLA-DR | APC Fire 810 | Biolegend |
| anti-human HLA-G | AlexaFluor 700 | Miltenyi |
| anti-human PD-1 | APC Cy7 | Biolegend |
| anti-human TIM3 | PE | Biolegend |
| anti-human CD14 | NovaFluor Yellow 610 | Biolegend |
| anti-human CD66b | PE/Fire 640 | Biolegend |
| anti-human CD117 | PE Cy5 | Biolegend |
| anti-human CD8 | NovaFluor Yellow 700 | Biolegend |
| anti-human PD-L1 | PE Cy7 | Biolegend |
| anti-human CD1c | BUV395 | Biolegend |
| anti-human CD206 | BUV737 | Biolegend |
| anti-human CD141 | BUV661 | Biolegend |

### Resource Availability

#### Lead Contact

Further information and requests for the resources and reagents should be directed to and will be fulfilled by the lead contact, Nina Bhardwaj.

#### Materials Availability

This study did not generate new unique reagents.

#### Data and Code Availability

We used raw RNAseq data from The Cancer Genome Atlas (TCGA) presented in two previous works and merged using TCGA identifiers ^13,59^. Data reported in this study are tabulated in the main text and supplementary materials.

Code is available on Github and GoogleColab.

https://github.com/gmestrallet/BasicScRNAseq

https://github.com/gmestrallet/BasicSpatialRNAseq

https://github.com/gmestrallet/IntegratingSpatialAndScRNAseq

https://github.com/gmestrallet/TCRscRNAseqAnalysis

Raw sequencing data are available on Mendeley.

https://data.mendeley.com/preview/ct2m5rfwvm?a=5b2c2c03-dcd9-474a-93b4-32642341d90c

#### Experimental model and subject details MSH2-deficient CT26, 4T1, B16F10 cells

CT26*, 4T1 and B16F10* cells deleted for MSH2 were provided by R.Samstein ^6^ and N.W.Cho ^60^. Guide RNA sequences (5’-CGGTGCGCCTCTTCGACCGC-3’) and (5’-GCACGGAGAGGACGCGCTGC-3’) targeting mouse Msh2 exon 1 were cloned into the PX461 plasmid and co-transfected into mouse CT26 mouse colon carcinoma cells using GenJetTM (SignaGen®) In Vitro DNA Transfection Reagent following the manufacturer’s protocol. 48 hours following transfection, GFP+ cells were seeded at one cell per 96-well by the Flow Cytometry Core Facility (FCCF). Cells were grown in RPMI supplemented with 10% FBS. Single cell clones were expanded and depletion of MSH2 was confirmed by Western Blot (anti-MSH2 monoclonal antibody (FE11), Invitrogen Antibodies). Confirmed Msh2-/-single cell subclones were expanded and used for serial passaging and downstream in vivo studies further described below.

## Method Detail

### Cell culture

Cells were amplified in RPMI media (Sigma) or DMEM media (Gibco) for cell culture. 10% FBS, 2 mM L-glutamine (Gibco) and 10% gentamicin were added to the medium. Medium was renewed 2 times a week. For cell amplification, cancer cells were seeded at 1,000 cells/cm2 and sub-cultured every week. All cultures were performed in plastic flasks (Biocoat, Becton-Dickinson). CT26 cells were serially passaged for continuous lengths of time under standard tissue culture conditions and were frozen under standard cell-specific freezing conditions with heat inactivated FBS and 10% DMSO cryoprotectant. Depletion of MSH2 was confirmed by Western Blot (anti-MSH2 monoclonal antibody (FE11), Invitrogen Antibodies).

### In vivo tumor growth analysis and ICB efficacy

2 * 10^5^ WT CT26 cells or CT26 MSH2 KO or 4T1 MSH2 KO or B16F10 MSH2 KO cells in 100 µl of PBS were orthotopically or subcutaneously injected in 6-week old BALB/c or BL6 mice (Jackson Laboratories). Mice with clinically palpable tumors (2 mm in diameter) were randomized into the following groups which received : isotype control IgG antibody, anti-mPD-1 Invivomab anti-mouse (CD279) (Bio X Cell), anti-TIM3 InVivoMab anti-mouse (CD366) (Bio X Cell), anti-mouse LAG-3 (Bio X Cell) (BE0174), anti-mouse CTLA-4 (CD152) (Bio X Cell), anti-mouse TREM2 (Leico), anti-mouse IL1B (Bio X Cell) or anti-TIGIT [1B4], Mouse IgG1, Lambda (Absolute Antibody) treatment groups (∼10 days post-injection). IgG or ICB antibodies (100µg) were administered intraperitoneally in 100 µl of PBS every 3-4 days (every 2 days for anti-TREM2, with a first injection of 200µg). Tumor volumes were measured weekly and calculated by the formula: (1/2) * (▭▭▭▭▭ℎ) * (▭▭▭▭ℎ)². 5 to 20 mice were used for each group. Mice were euthanized by carbon dioxide and the tumors were resected 28 days after tumor injection.

### TCGA MSI and CIBERSORT analysis

We used raw RNAseq data from The Cancer Genome Atlas (TCGA) presented in two previous works and merged using TCGA identifiers ^13,59^. Patient MSI status was assessed using the MSI sensor. Patient MSI status was considered as MSI-H for a MSIsensor threshold of 3.5. MSI-H patients of Uterine Corpus Endometrial Carcinoma (UCEC) and Colon Adenocarcinoma (COAD) cohorts were considered for CIBERSORT (cell-type identification by estimating relative subsets of RNA transcripts) analysis. The relative fraction of 22 immune cell types within the leukocyte compartment could be estimated using CIBERSORT ^61^. Linear regressions between various immune parameters were performed using Python.

### Spheroid growth

10,000 tumor cells were seeded in 200µL of RPMI medium in a nucleon sphera 96 well plate (Thermofisher). As an alternative, spheroids were grown using SpheroTribe (Idylle) ^62^. After 1 to 3 days, the circularity and the area of spheroids were measured as previously described in other systems ^63^. Circularity = (4π Area) / (perimeter)², Roundness = 4 Area / (▭ × major axis²), with Area in µm². All parameters were measured using the Image J software.

### Spheroid infiltration by PBMC and ICB efficacy

10,000 cells were seeded in 96 low attachment well plates (Nucleonsphera, Thermofisher) with RPMI medium, to form spheroids. PBMC were seeded in other 96-well culture plates (Thermofisher) and incubated at 37 °C, in 5% CO2 in 200 μL RPMI medium supplemented with 20% FCS, enriched in gentamicin. 100,000 PBMCs were activated per well by IL-15 (40ng/mL) and treated or not with ICB (10µg/mL of anti-PD-1, anti-LAG3, anti-TIM3 or anti-TIGIT) for 3 days. Then, these 100,000 PBMC were added to each tumor spheroid (10,000 cells) for 3 days. Spheroids incubated with PBMC were isolated from PBMCs in suspension and dissociated using accutase (Corning Media Tech) with 3 cycles of 10 minutes or using MACS dissociator and a mouse tumor dissociation kit (Miltenyi). Then, PBMC, spheroids or spheroids incubated with activated PBMC were labeled with antibodies listed in **Table 1** and Live Dead Blue (1uL of each antibody in 200uL of PBS). Collection was made using an Attune NxT flow cytometer (Thermofisher) or a spectral Aurora (Cytek). Data were analyzed using FlowJo software (BD biosciences). PBMC ability to infiltrate spheroids was analyzed by comparing CD45 expression on spheroids incubated with PBMC in different conditions. Tumor cell death was measured using propidium iodide (5μg/mL).

### Flow cytometry analysis

Fresh tumors were resected and dissociated into single cell suspensions using a gentle MACS tissue dissociator and mouse tumor dissociation kit (Miltenyi). For analysis of cell-surface immune markers expression, cancer cells were processed as single-cell suspensions and stained for 30 minutes at room temperature with monoclonal antibodies. PBMC were labeled with antibodies listed in **Table 1** and Live Dead Blue (1uL of each antibody in 200uL of PBS). Non-reactive antibodies of similar species and isotype, and coupled with same fluorochromes, were used as isotypic controls. Immune expression profiles were analyzed using an Attune NxT (Thermofisher) or a spectral Aurora (Cytek). Data were analyzed using FlowJo software (BD biosciences).

### Bi-photon spheroid imaging

CT26 spheroids deleted for MSH2 were grown for 3 days and then co-cultured with or without immune cells for 3 more days and ICB. Splenocytes immune cells are previously labeled by Life Technologies celltracker Deep Red (Thermo Scientific). Spheroid cell nuclei are labeled with Molecular Probes NucBlue Fixed Cell ReadyProbes Reagent (Thermo Scientific). 3D Bi-photon imaging of spheroid infiltration by immune cells was performed using Olympus FVMPE-RS. Spheroid immune infiltration was followed on the (x/y) axis in the z-middle. Z-stack imaging of spheroid immune infiltration (top to middle of the spheroid) was also performed.

### Whole Exome Sequencing (WES)

A sample sheet was generated to document murine sample details, including sample paths, using a format specifying murine ID, sample type (normal or tumor), lane information, and corresponding FASTQ files. The sample sheet, named “samplesheet.csv,” was organized within the directory containing the associated FASTQ files. A job file, denoted as “submit_job.sh,” was created to facilitate variant calling using Nextflow. This job file incorporated necessary module additions for Java (version 11.0.2) and Singularity (version 3.2.1). The Nextflow script executed the variant calling pipeline from nf-core/sarek, tailored for WES data analysis ^64,65^. Input parameters included the path to the sample sheet (--input), maximum CPU allocation (--max_cpus), specification for only paired variant calling (--only_paired_variant_calling), selected variant calling tools (e.g., Strelka, Mutect2), reference genome (e.g., GRCm38), and output directory (--outdir). Nextflow was set up by installing the required modules for Java (version 11.0.2) and Singularity (version 3.2.1). The Nextflow executable was downloaded and configured in the user’s home directory (∼/nextflow). Verification of the Nextflow installation and version was performed to ensure proper functionality. Upon setup completion, variant calling jobs were initiated for each sample using the provided job file (submit_job.sh). These jobs were submitted to the computing cluster for execution, with subsequent monitoring until completion. Variant annotation was conducted using Python scripting with the Varcode library. Variant Call Format (VCF) files were loaded, correcting genomic mismatches as necessary.

### Single cell, TCRseq and spatial RNAseq analysis

8 CT26 WT and 8 MSH2 KO tumors were inoculated for 4 weeks in BALC mice as described above. Anti-PD-1 was administered twice a week after the 2 first weeks in half of the mice. Tumors were harvested after 4 weeks, and 4 tumors (1 per condition) were fixed in PFA and ethanol, and incorporated in paraffin blocks to be sent to spatial RNA sequencing by the human immune monitoring group at Mount Sinai. 12 tumors (3 per condition) were sent for single cell RNA sequencing by the human immune monitoring group. Downstream analysis of 63807 cells was performed using python and scanpy. All samples were preprocessed to remove genes that are expressed in less than 3 cells and to identify mitochondrial genes. Cells that have too many mitochondrial genes were removed from the analysis. Data were normalized and highly-variable genes were identified. Principal component analysis was performed and clustering was performed using the leiden graph-clustering method. Marker genes were identified for each cluster and a secondary clustering was performed to separate the immune cells and the cancer cells with the Wilcoxon Rank-Sum test. Gene expression was analyzed in the different conditions using scanpy. TCRseq was performed using Scirpy. Single cell clustering was finally applied to the spatial samples using scanorama and scanpy. A RandomForestClassifier algorithm was also trained on a scRNAseq dataset ^17^, n=10 MMRd CRC patients. Accuracy = correct predictions / total number of predictions. Precision = correct predictions of a class / all positive true positive and false positive predictions. Recall (Sensitivity) = correct true positive predictions of a class / actual instances of the class. F1-score = harmonic mean of precision and recall.

### Data analysis and visualization

Spectral flow cytometry and RNAseq data were analyzed using Python 3 and Jupyter Notebook. Numpy and Pandas were used for array and data frame operations, and data visualization was performed using Matplotlib and Seaborn. scRNA-seq data were analyzed using Cell Ranger and Scanpy. Flow cytometry data were analyzed using FlowJo and Python 3.

### Statistics

Statistical significance of the observed differences was determined using the Mann-Whitney two-sided U-test, T-test or Linear regression. All data are presented as mean±SEM. The difference was considered as significant when the p value was below 0.05. * : p<0.05, ** : p<0.01 and *** : p<0.001.

## Acknowledgments

The authors thank all the current and previous members of the Bhardwaj, Samstein, Merad and Vabret labs for critical comments on the manuscript. The authors also thank the members of the Human Immune Monitoring Center for processing the Luminex, scRNAseq and spatial RNAseq samples and providing the data, the members of the flow cytometry core, the microscopy core and the pathology core at Mount Sinai Hospital for helpful discussions and training to prepare the experiments. This work was supported in part through the computational and data resources and staff expertise provided by Scientific Computing and Data at the Icahn School of Medicine at Mount Sinai and supported by the Clinical and Translational Science Awards (CTSA) grant UL1TR004419 from the National Center for Advancing Translational Sciences. Research reported in this publication was also supported by the Office of Research Infrastructure of the National Institutes of Health under award number S10OD026880 and S10OD030463. The content is solely the responsibility of the authors and does not necessarily represent the official views of the National Institutes of Health.

## Author Contributions

G.M., C.C.B., R.S. and N.B. conceptualized the project. G.M. performed the in vivo and in vitro experiments. G.M. performed the analysis of the genetic, flow cytometry and microscopy data. M.B. helped with WES analysis. N.Vaninov helped with the in vivo orthotopic experiments for the 4T1 model. L.V. and A.A. provided human blood samples and other reagents. N.W.C. and M.S. provided the polyclonal CT26 MMRd cell line and financed WES. G.M. wrote the manuscript. G.M., N.Vabret, C.C.B., R.S. and N.B. revised the manuscript.

## Supplementary data

**Table 1.**
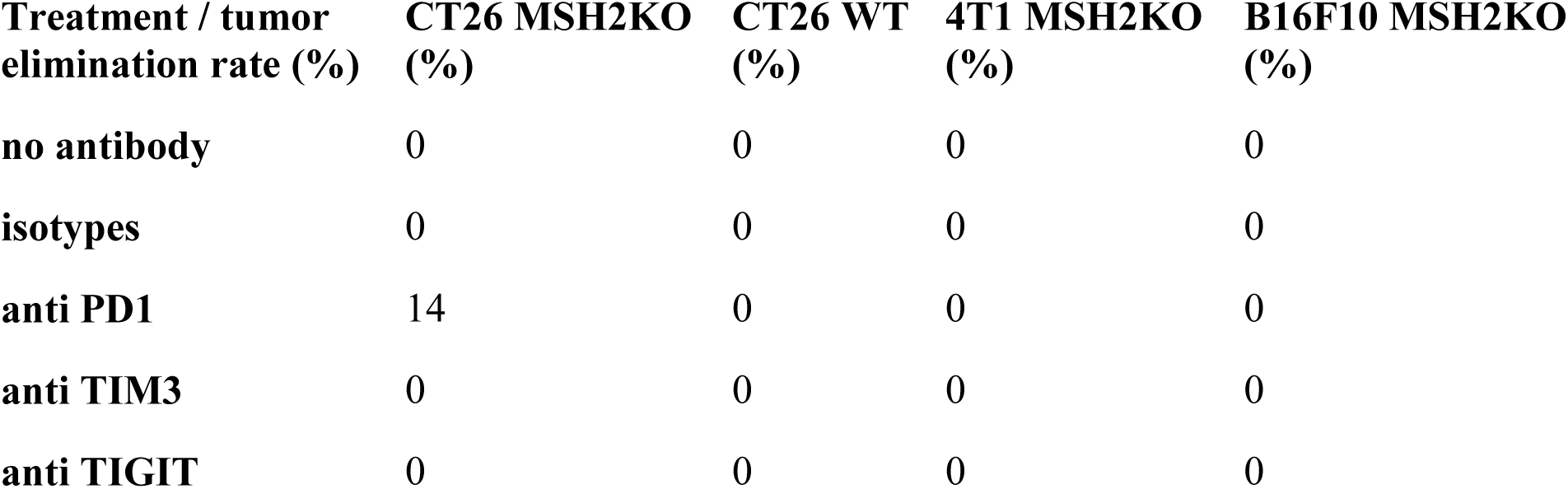
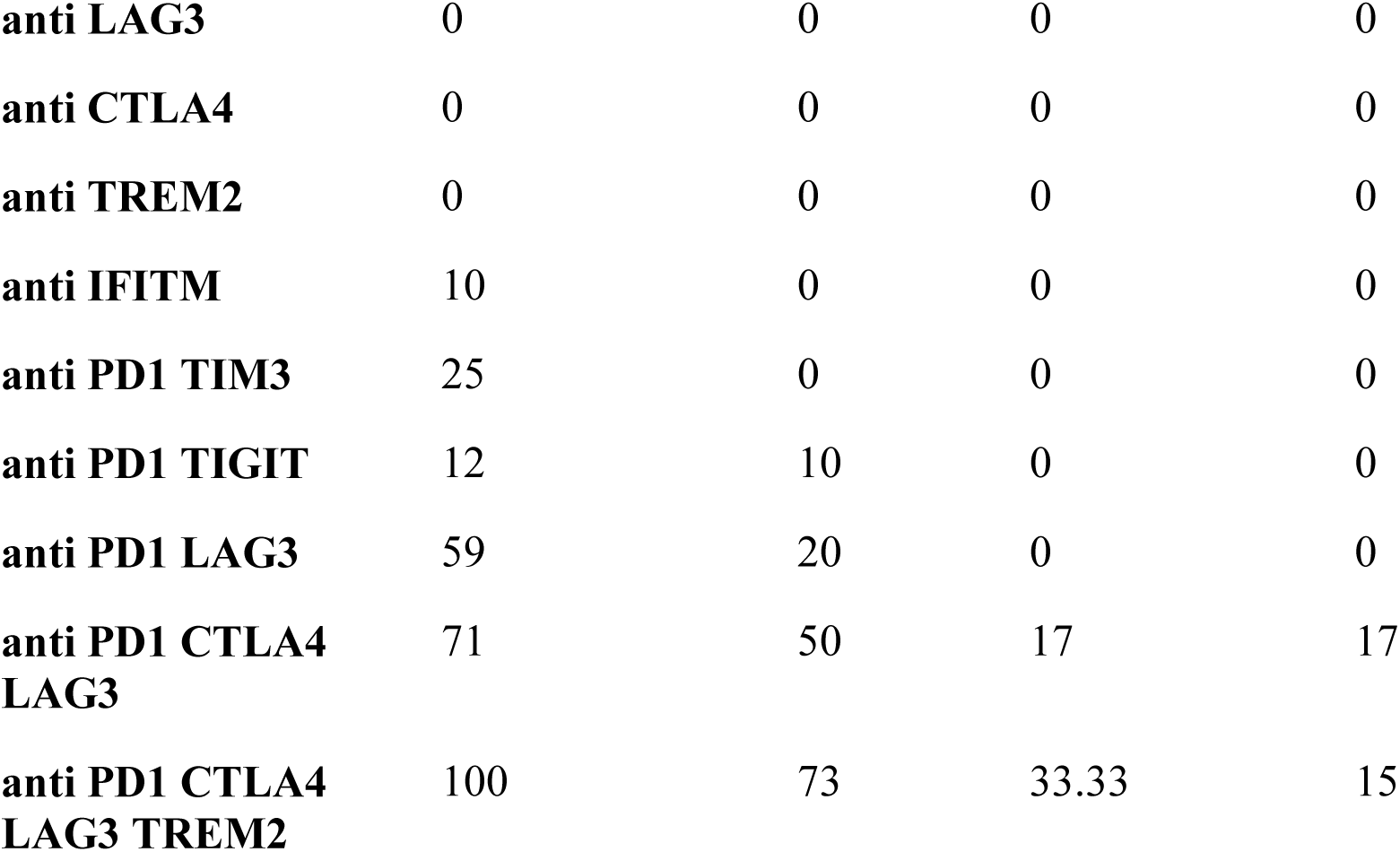
Percentage of complete tumor elimination in mice according to treatment.

| **Treatment / tumor elimination rate (%)** | **CT26 MSH2KO (%)** | **CT26 WT (%)** | **4T1 MSH2KO (%)** | **B16F10 MSH2KO (%)** |
| --- | --- | --- | --- | --- |
| **no antibody** | 0 | 0 | 0 | 0 |
| **isotypes** | 0 | 0 | 0 | 0 |
| **anti PD1** | 14 | 0 | 0 | 0 |
| **anti TIM3** | 0 | 0 | 0 | 0 |
| **anti TIGIT** | 0 | 0 | 0 | 0 |
| **anti LAG3** | 0 | 0 | 0 | 0 |
| **anti CTLA4** | 0 | 0 | 0 | 0 |
| **anti TREM2** | 0 | 0 | 0 | 0 |
| **anti IFITM** | 10 | 0 | 0 | 0 |
| **anti PD1 TIM3** | 25 | 0 | 0 | 0 |
| **anti PD1 TIGIT** | 12 | 10 | 0 | 0 |
| **anti PD1 LAG3** | 59 | 20 | 0 | 0 |
| **anti PD1 CTLA4**<br>**LAG3** | 71 | 50 | 17 | 17 |
| **anti PD1 CTLA4**<br>**LAG3 TREM2** | 100 | 73 | 33.33 | 15 |

**Extended Figure 1.**
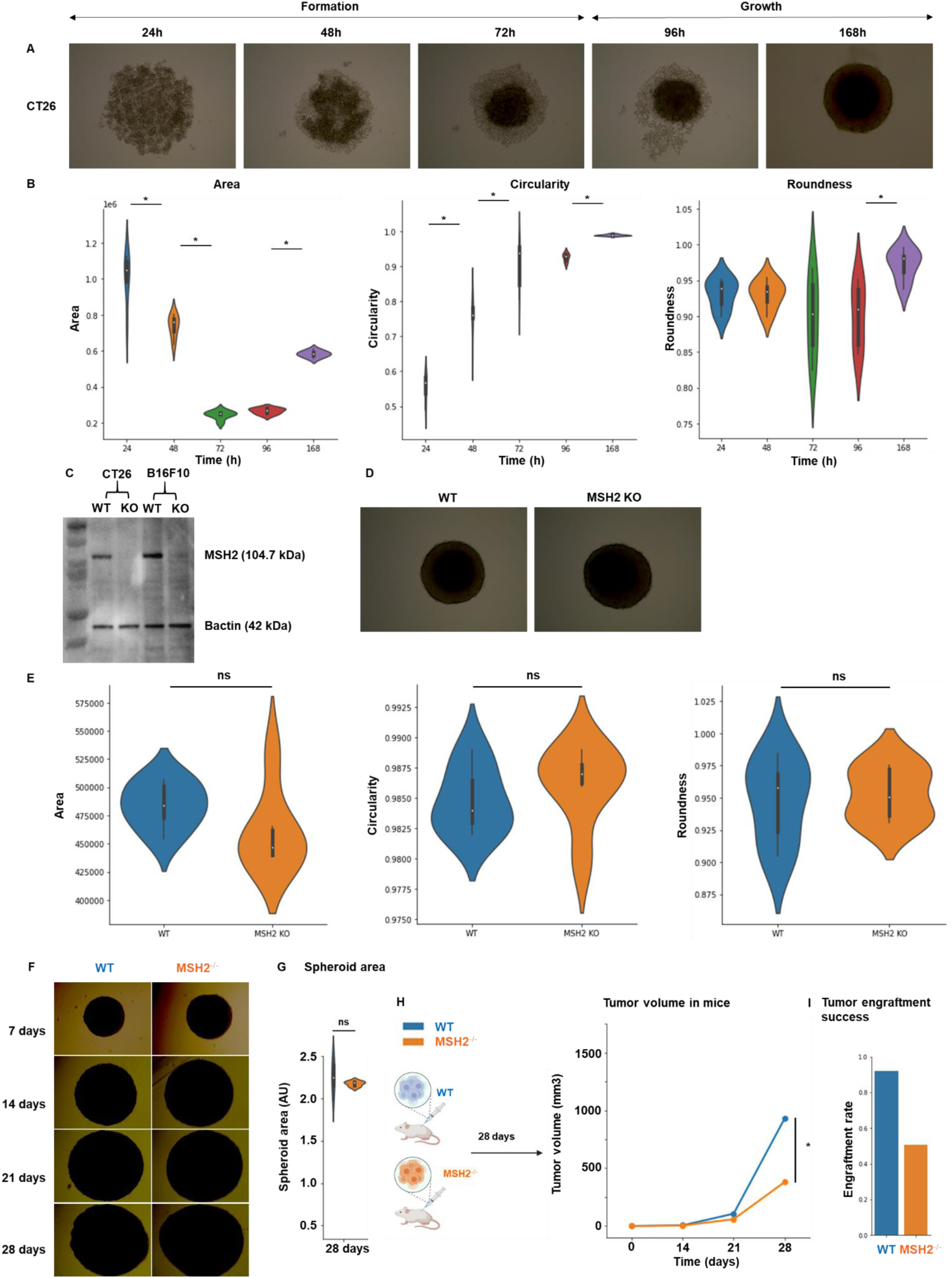
Effect of the TME on MSH2 KO tumor development. **A** 10,000 WT CT26 thawed cells were used per spheroid in a 96 well plate. **B** After 1,2,3,4 and 7 days, roundness, circularity and area of CT26 spheroids were measured using ImageJ software. N=6 replicates. **C** MSH2 was deleted in CT26 and B16F10 cells using Crispr and cell lines were subcloned. Western Blot to assess MSH2 deletion was performed using anti-Bactin antibody (Abcam, clone 8226), anti MSH2 antibody (Thermo Scientific, clone FE11) and anti-mouse secondary antibody (Thermo Scientific, clone A28177). N=3 **D** 10,000 WT or MSH2 KO CT26 cells were used per spheroid in a 96 well plate. **E** After 3 days, roundness, circularity and area were measured using ImageJ software. N=6 replicates. **F** 10,000 CT26 cells were used per spheroid in a 96 well plate. **G** After 28 days, the area was measured using ImageJ software. N=6 replicates. **H** 200,000 CT26 tumors, either deleted or not for MSH2, were allowed to grow for 28 days in BALB/C mice. Tumor volume during growth (in mm3). N= 7 to 27. **I** Tumor engraftment rate (in %). N= 26 to 48. Mann-Whitney U Test was used for all comparisons, and the difference was deemed significant if p value<0.05.

**Extended Figure 2.**
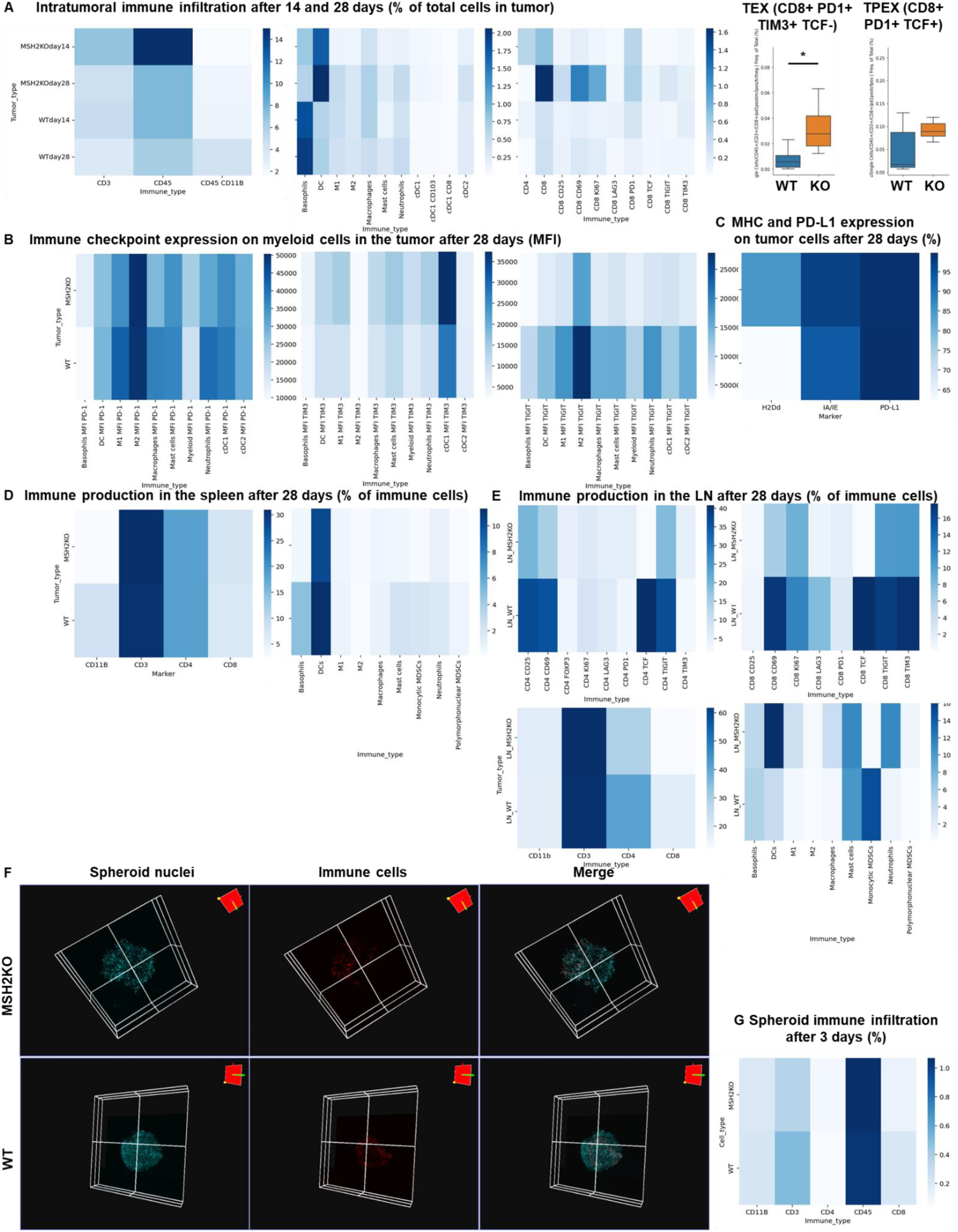
Immune landscape of MSH2 KO and WT tumors and spheroids, spleen and lymph nodes. CT26 tumors deleted or not for MSH2 are growing for 28 days in BALB/C mice. **A** Quantification of differential immune checkpoint expression and immune infiltration according to the MSI status after 14 days and 28 days. N=6 replicates, Mann-Whitney U Test, p value<0.05 **B** Immune checkpoint expression on tumor infiltrating myeloid cells after 28 days. **C** MHC and PD-L1 expression by tumor cells after 28 days. **D/E** Immune production in the spleen/lymph nodes of mice injected by WT and MSH2 KO tumors after 28 days. **F** CT26 spheroid deleted for MSH2 are growing for 28 co-cultured with immune cells for 3 days. Splenocytes immune cells are previously labeled by cell tracker (red). Spheroid cell nuclei are labeled with Nunc blue (blue). 3D Bi-photon imaging. **G** 10,000 CT26 cells deleted or not for MSH2 are growing for 3 days as spheroids and 100,000 PBMC from 3 spleens are activated with IL-15. Then, spheroids and PBMC are incubated for 3 days. Flow cytometry staining of spheroids after dissociation. Quantification of differential immune checkpoint expression and lymphocyte infiltration according to the MSH2 status. N=6. Mann-Whitney U Test, p value<0.05

**Extended Figure 3.**
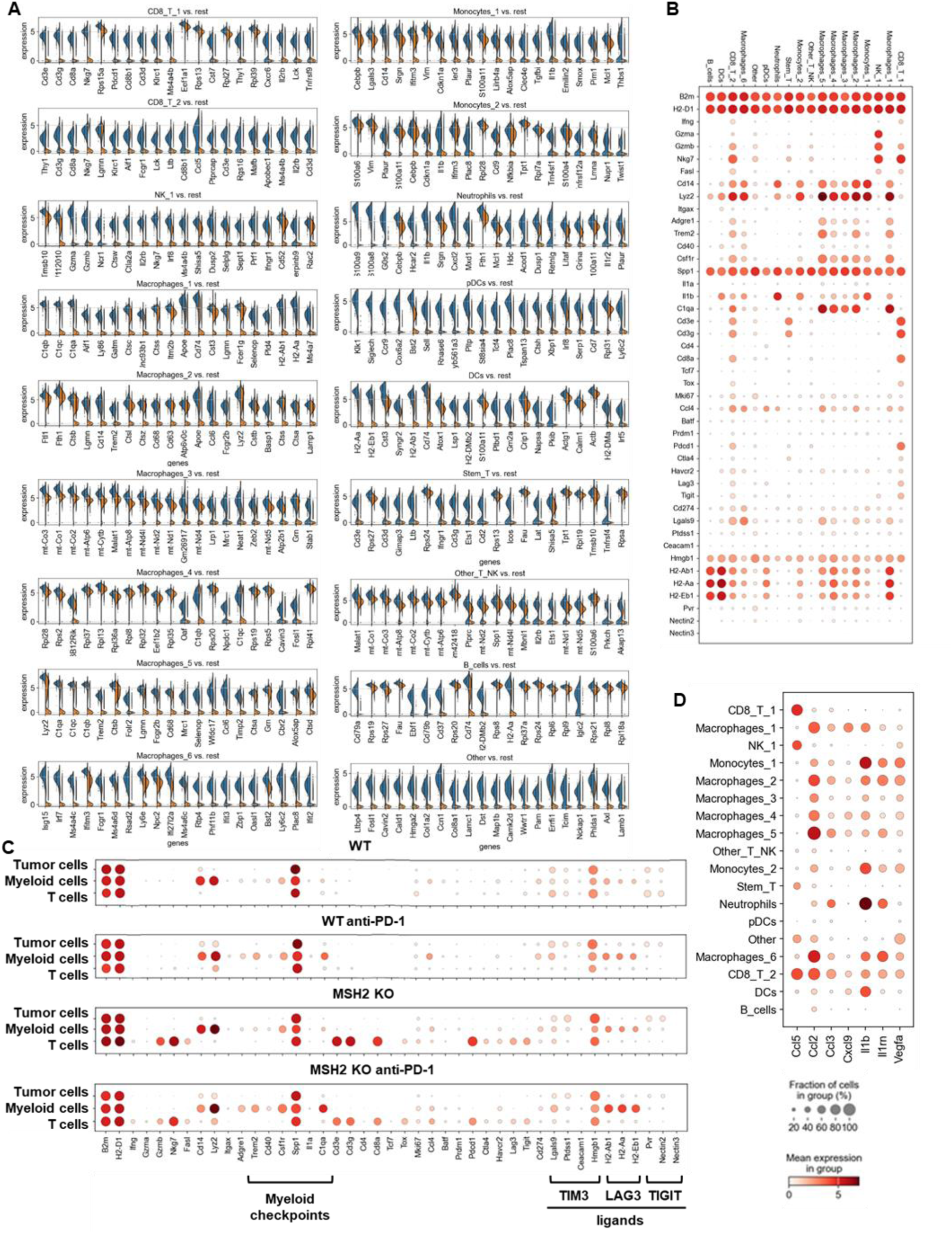
Single cell analysis of immune cells in CT26 WT and MSH2 KO tumor with or without anti-PD-1 therapy. ScRNASeq of immune cells in CT26 WT and MSH2 KO tumor with or without anti-PD-1 therapy. **A** More significantly upregulated genes in each immune cluster (Wilcoxon test). **B** Immune checkpoint and myeloid immunosuppressive molecule expression in each immune cluster. **C** Single cell analysis of WT and MSH2 KO tumor and immune cells after 28 days, with or without anti-PD-1 therapy. **D** cytokine production by immune cells. N=3 tumors per condition.

**Extended Figure 4.**
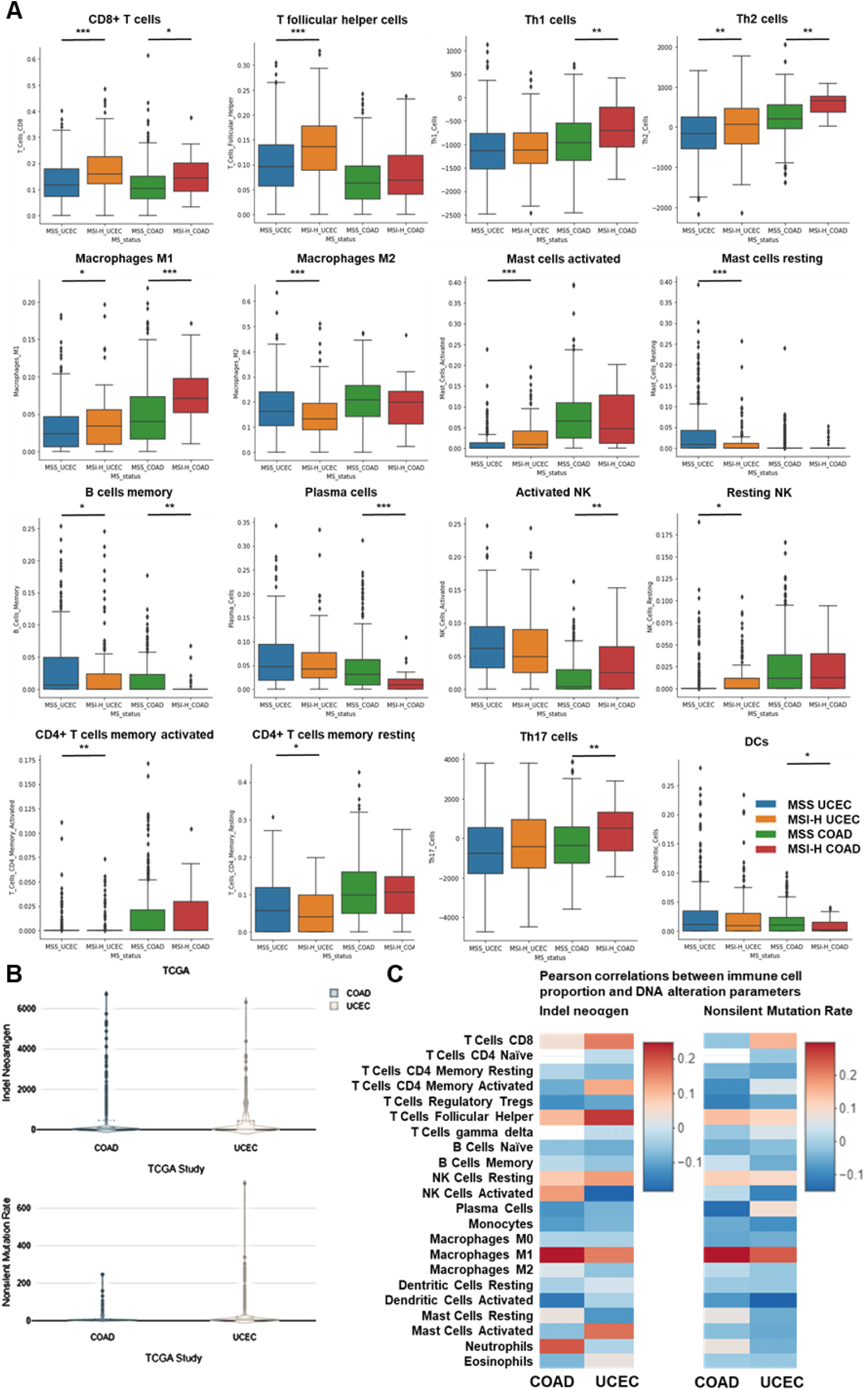
Immune infiltration in MMRd UCEC and COAD human tumors. **A** CIBERSORT TCGA analysis of UCEC (N = 528 patients) and COAD (N = 442 patients) tumor infiltration by immune cells. Mann-Whitney U Test, p value<0.05 **B** Indel neoantigen load and Non-silent mutation rate in COAD and UCEC TCGA cohorts on CRI iAtlas. **C** Pearson correlations between immune cell proportion and DNA alteration parameters in CRI iAtlas TCGA UCEC (N = 543 patients) and COAD (N = 445 patients) cohort.

**Extended Figure 5.**
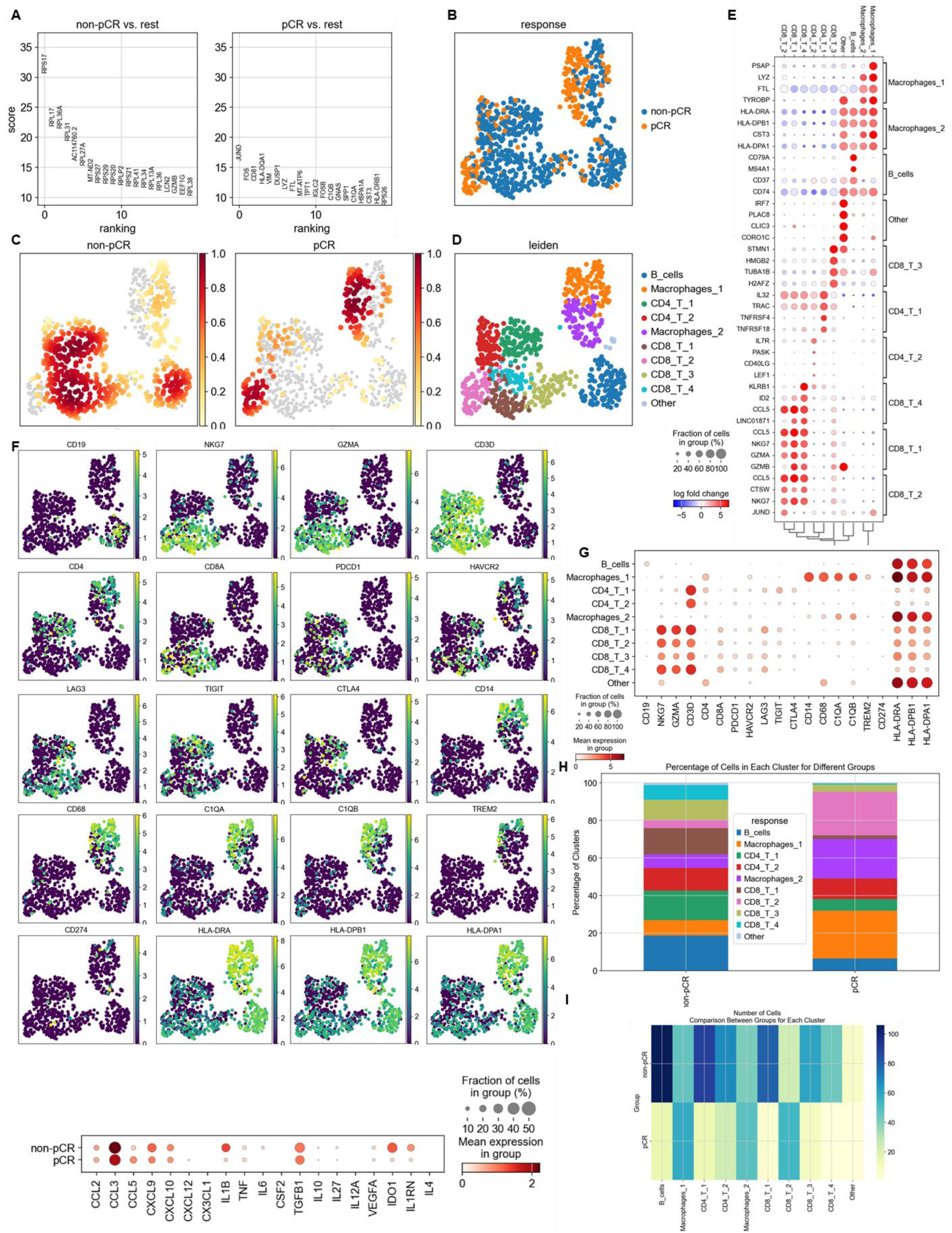
Single cell RNAseq analysis of immune cells in MMRd CRC patients that responded or not to anti-PD-1 therapy. We reanalyzed human MMRd CRC patient data previously published [Jianxia Li et al, Cancer cell 2023] with a focus on patients that completely responded or not to anti-PD-1 therapy and the underlying T cell and macrophage profiles. **A** More significantly upregulated genes in immune cells in patients that responded or not to anti-PD-1 therapy (Wilcoxon test). **B** ScRNAseq UMAP of immune cells in patients that responded or not to anti-PD-1 therapy. **C** Density UMAP of immune cells in patients that responded or not to anti-PD-1 therapy. **D** UMAP of immune clusters following leiden clustering by scanpy. **E** More significantly upregulated genes in each immune cluster. **F/G** Immune checkpoint and myeloid immunosuppressive molecule expression in each immune cluster. **H/I** Percentage/number of cells in each immune cluster.

**Extended Figure 6.**
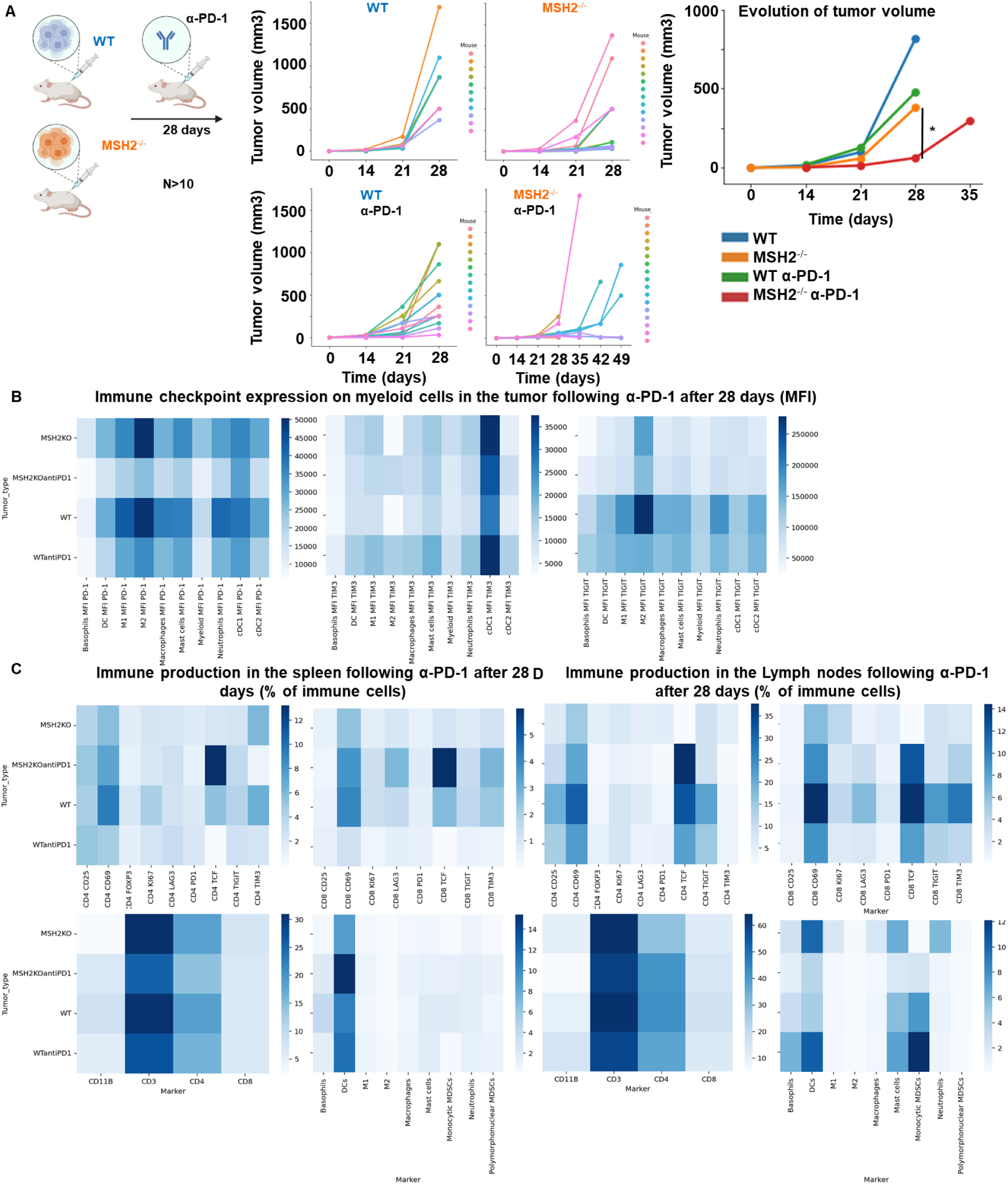
Immune landscape of MSH2 KO and WT tumors, spleen and lymph nodes after anti-PD-1 therapy. **A** CT26 tumors deleted or not for MSH2 are growing for 28 days in BALB/C mice, with or without anti PD-1 twice a week after 14 days. **B** Immune checkpoint expression on tumor infiltrating myeloid cells after 28 days following anti-PD-1. **C/D** Immune production in the spleen/lymph nodes of mice injected by WT and MSH2 KO tumors after 28 days following anti-PD-1. N=6 replicates, Mann-Whitney U Test, p value<0.05

**Extended Figure 7.**
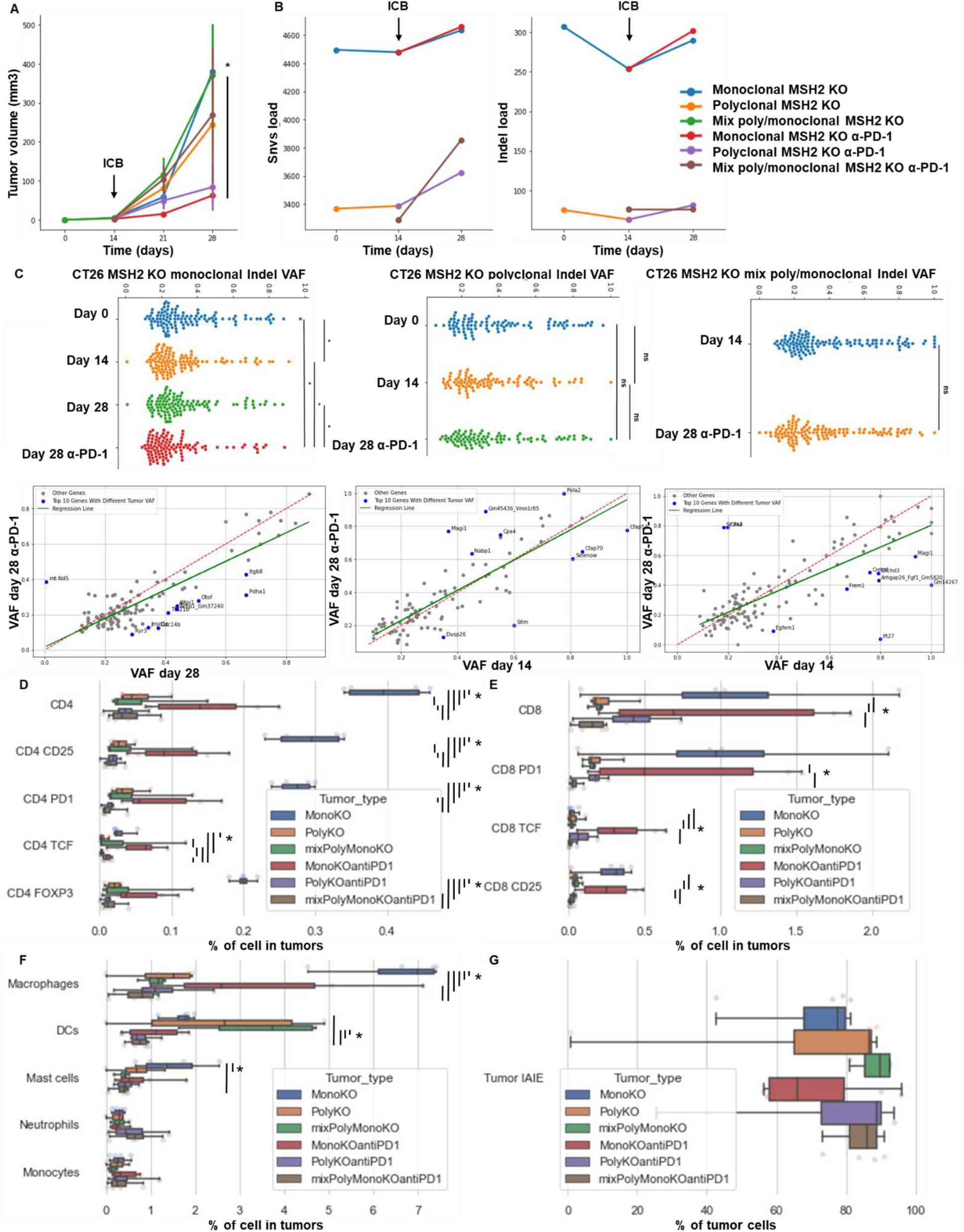
High indel and snv loads and infiltration by TCF+ CD4+ and CD8+ T cells correlate with longitudinal response to ICB and immunoediting. Monoclonal or polyclonal CT26 tumors deleted for MSH2 are growing for 28 days in BALB/C mice, with or without anti PD-1 twice a week after 14 days. **A** Tumor volume in vivo over 28 days in the absence or presence of anti-PD-1 (100 µg twice a week after 14 days) therapy (in mm3). N= 6 to 27, Mann-Whitney U Test, pvalue<0.05. **B** Indel and snv count with effect on the coding sequence following whole exome sequencing (strelka, varcode). **C** Quantification of tumor mutational allele frequency changes following ICB. Mann-Whitney U Test, pvalue<0.05. Quantification of **D** CD4 + T cell, **E** CD8+ T cell and **F** myeloid infiltration according to the MSI status and ICB. **G** Quantification of MHCII expression on tumors. N=4 replicates, Mann-Whitney U Test, pvalue<0.05

**Extended Figure 8.**
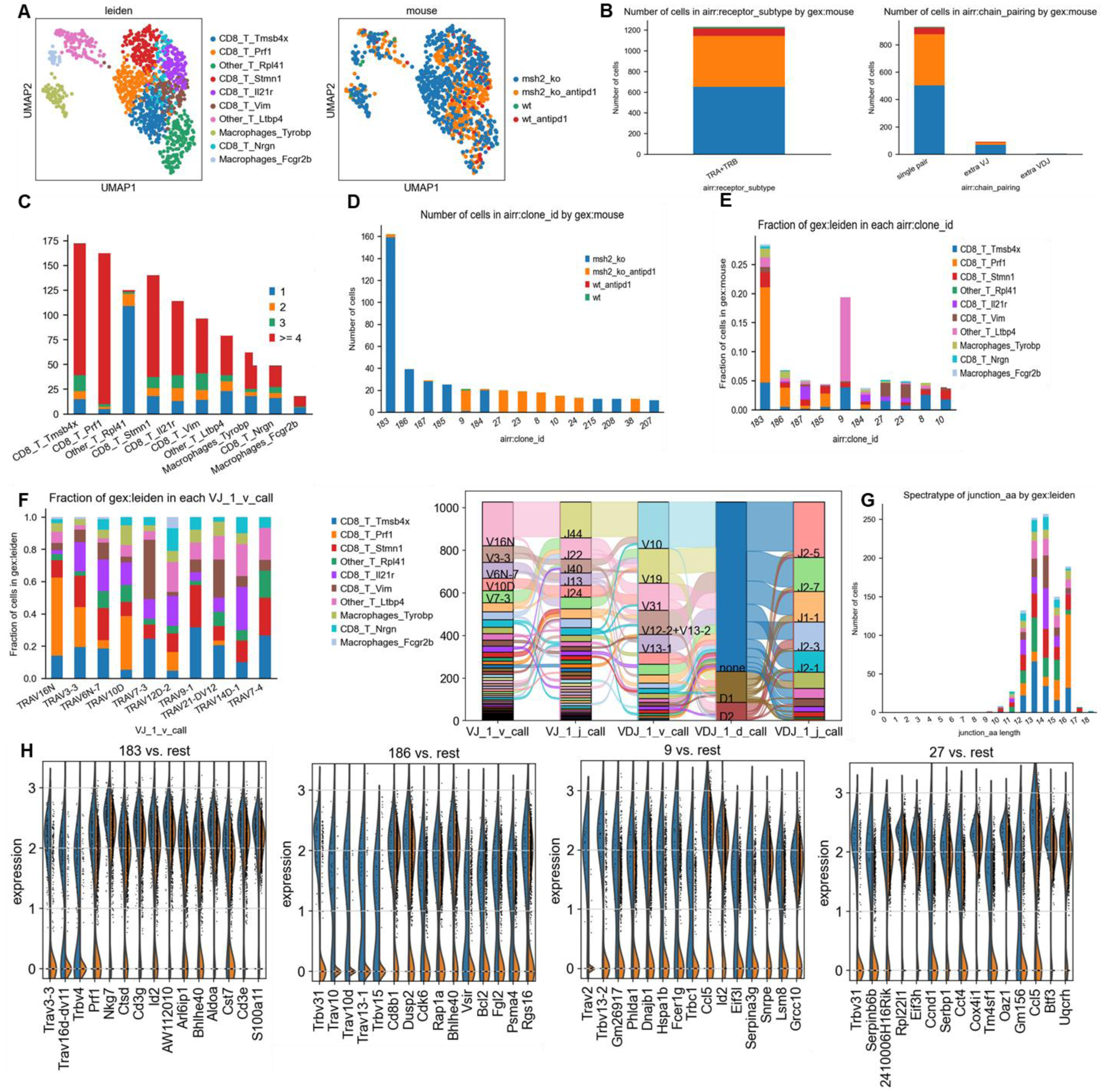
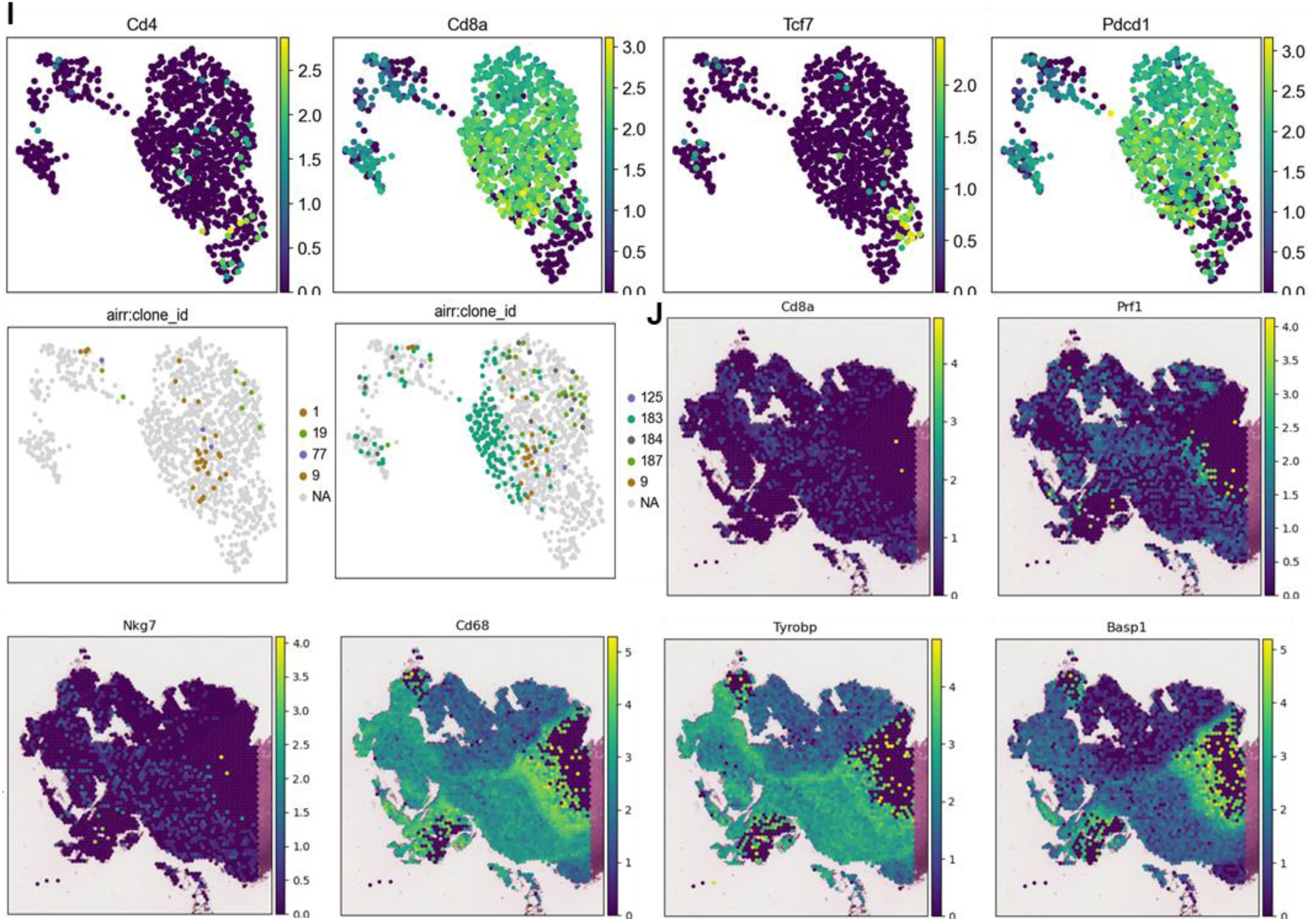
TCRseq analysis paired with scRNAseq analysis on MMRd tumor according to anti-PD-1 therapy. **A** scRNAseq clustering on immune cells infiltrating tumors expressing TCRs. **B** The dataset contains only α/β T-cell receptors. Filtering to exclude all cells that don’t have at least one full pair of receptor sequences and multichain cells. **C** Clonal expansion, number of cells in each clonotype. **D/E** Clonotype abundance in each cluster and in each condition. **F** 10 most abundant V-genes and exact combinations of VDJ genes. **G** Length distribution of CDR3 regions. **H** Overexpressed genes in the more abundant clonotypes. **I** Gene expression in clones shared between different tumor types. **J** Spatial localization of main CD8 and macrophage clones in MSH2 KO tumors after ICB.

**Extended Figure 9.**
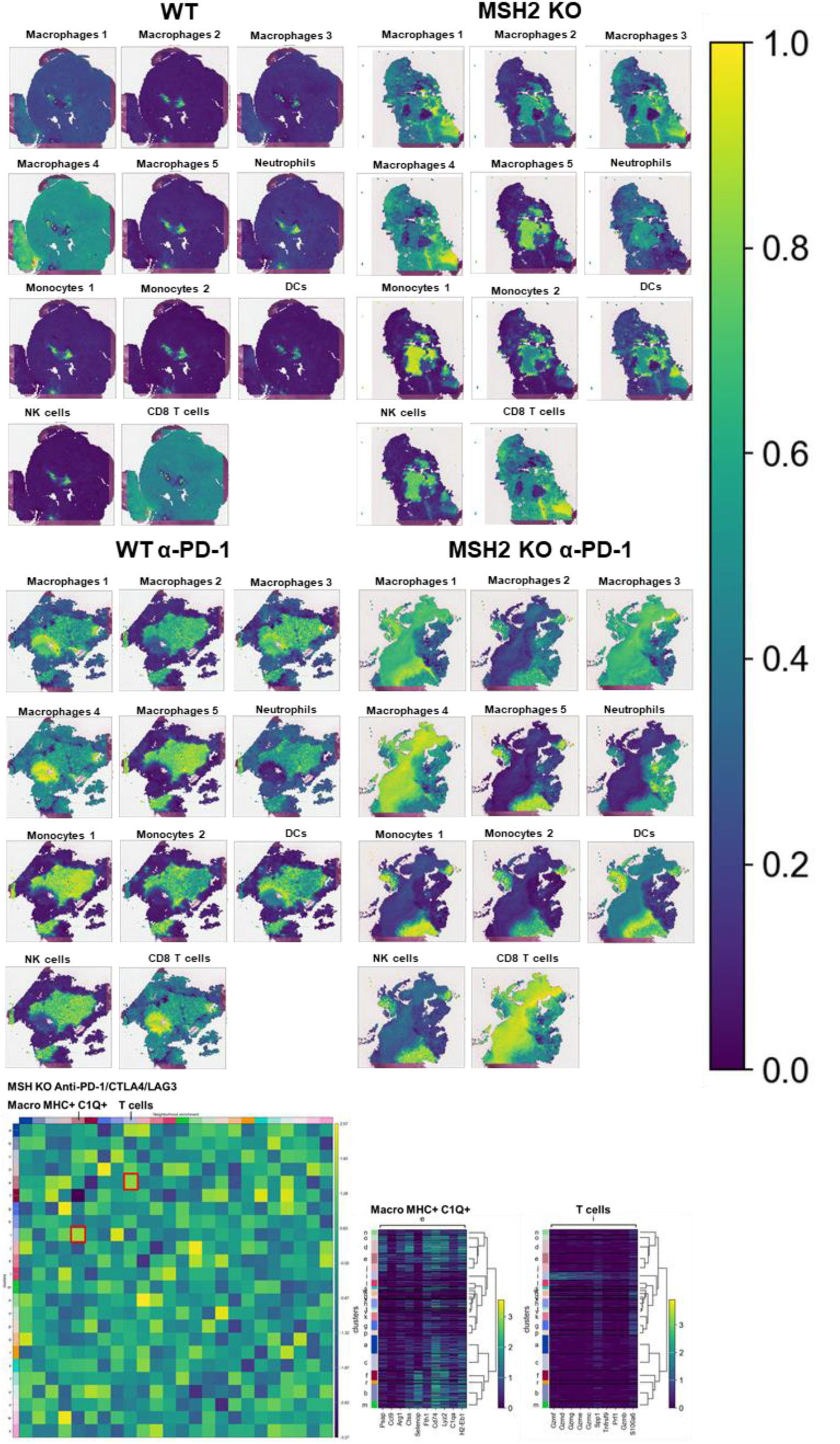
Spatial RNAseq analysis of immune cells in CT26 WT and MSH2 KO tumor with or without ICB therapy. Spatial localization of each scRNAseq immune cluster in CT26 WT and MSH2 KO tumor with or without ICB therapy. Neighborhood enrichment was performed using scanpy and squidpy.

**Extended Figure 10.**
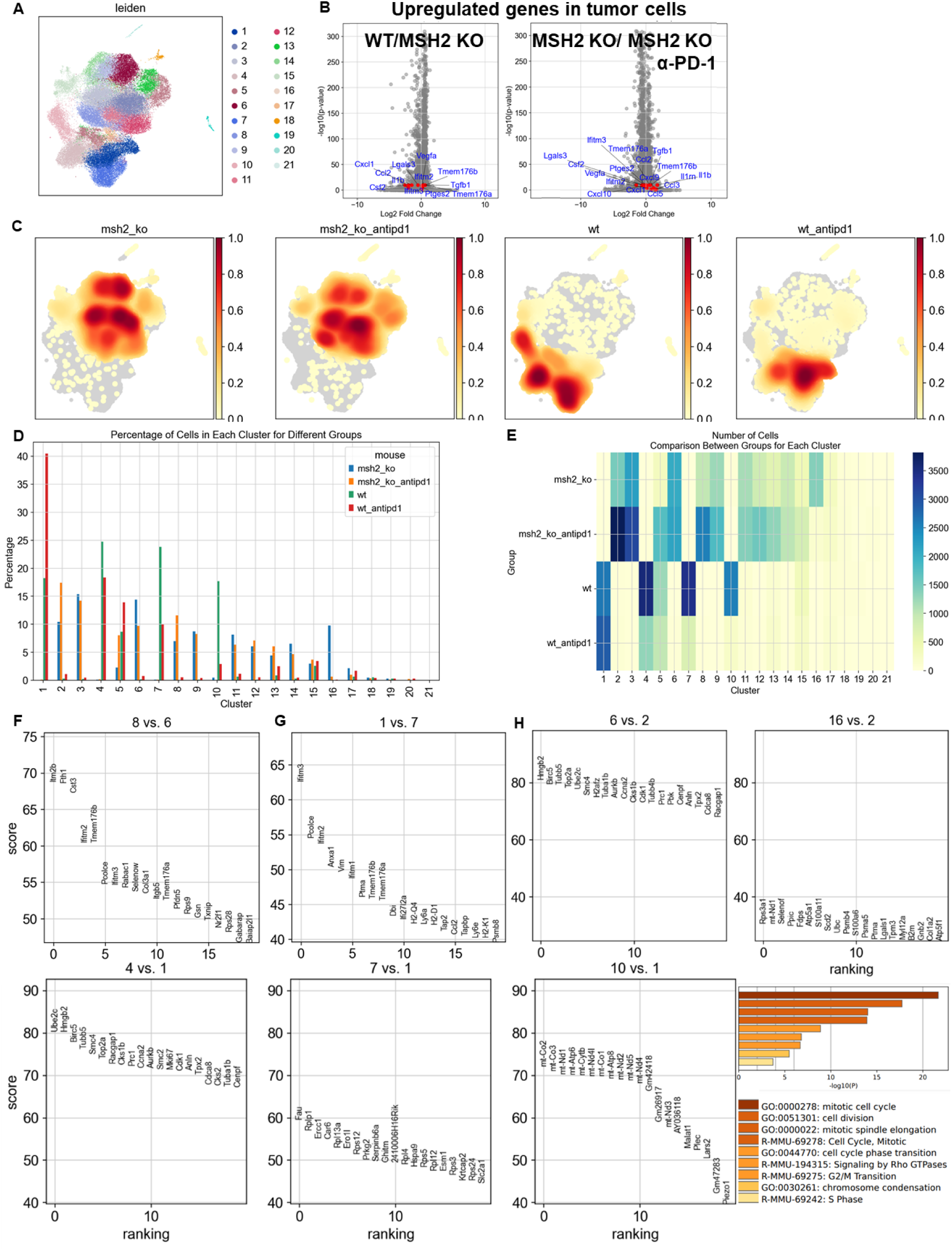
Single cell RNAseq analysis of tumor cells in CT26 WT and MSH2 KO tumor with or without anti-PD-1 therapy. **A** ScRNAseq UMAP of tumor cells, after performing exclusion of immune cells, in CT26 WT and MSH2 KO tumor with or without anti-PD-1 therapy, following leiden clustering by scanpy. **B** More significantly upregulated genes in tumor cells in CT26 WT and MSH2 KO tumor with or without anti-PD-1 therapy (Wilcoxon test). **C** Density UMAP of tumor cells in CT26 WT and MSH2 KO tumor with or without anti-PD-1 therapy. **D/E** Percentage/number of cells in each tumor cluster. **F** More significantly upregulated genes in MSH2 KO tumor cluster resisting anti-PD-1 therapy compared to cluster responding to anti-PD-1 (Wilcoxon test). **G** More significantly upregulated genes in WT tumor cluster resisting anti-PD-1 therapy compared to cluster responding to anti-PD-1 (Wilcoxon test). **H** More significantly upregulated genes in WT and MSH2 KO tumor clusters targeted by anti-PD-1 therapy compared to cluster resisting anti-PD-1 (Wilcoxon test) and corresponding gene enrichment analysis with metascape.

**Extended Figure 11.**
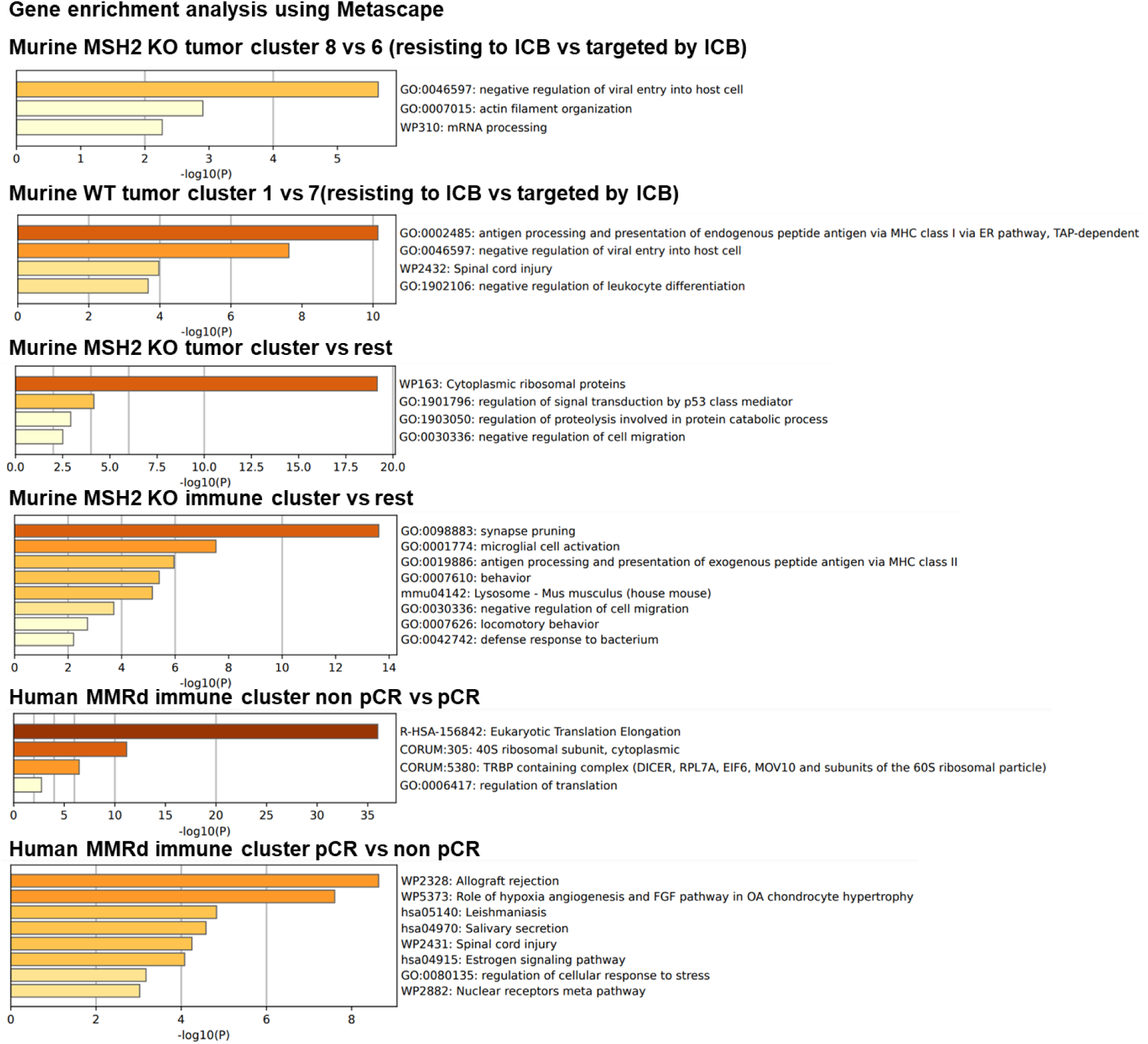
Gene enrichment analysis in immune and tumor scRNAseq clusters. Gene enrichment analysis was performed using Metascape ^66^ in immune and tumor scRNAseq clusters.

**Extended Figure 12.**
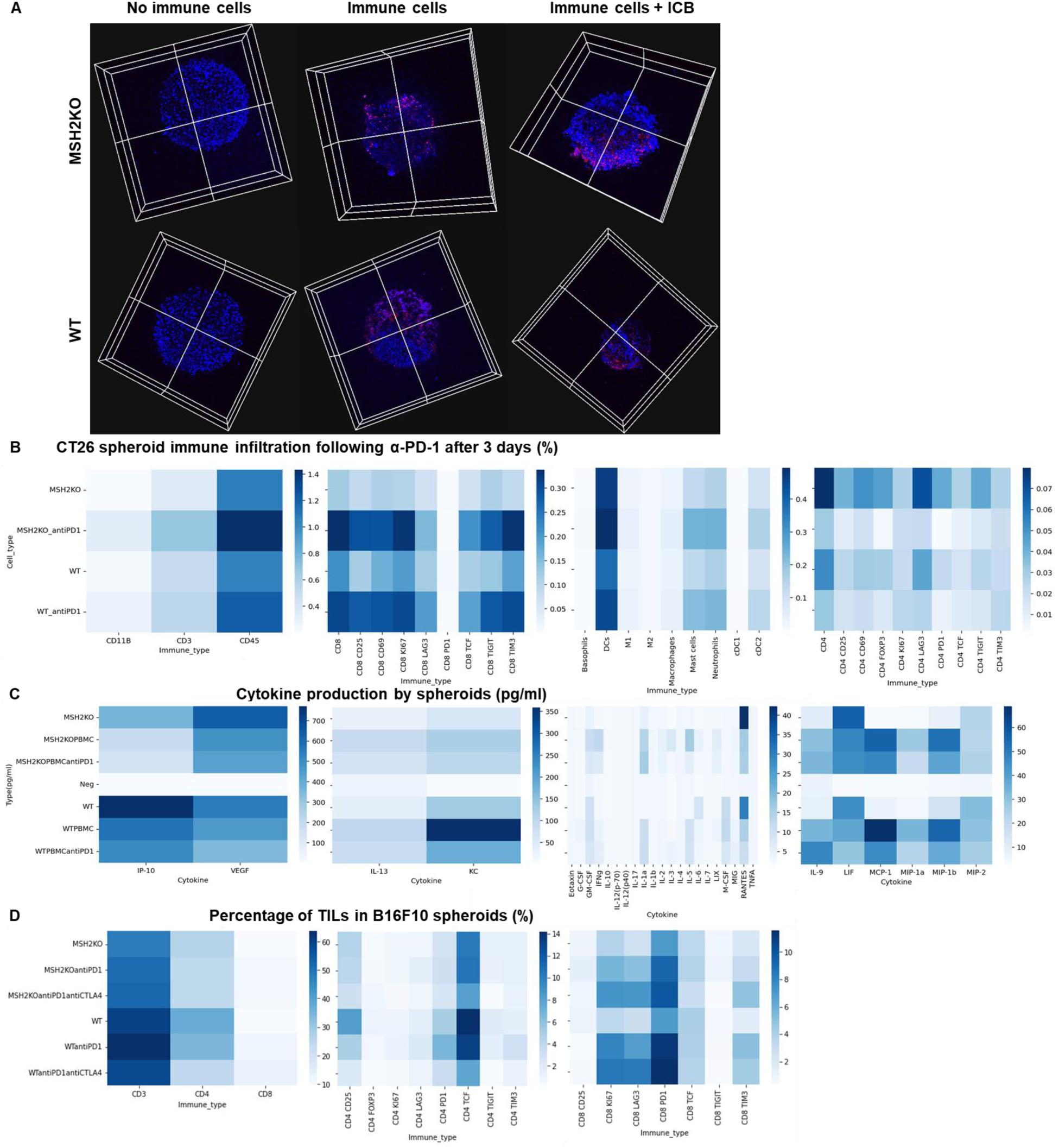
Immune landscape of MSH2 KO and WT spheroids after anti-PD-1 therapy. **A** CT26 spheroids deleted for MSH2 are growing for 3 days and then co-cultured with or without immune cells for 3 more days and ICB. Splenocytes immune cells are previously labeled by cell tracker (red). Spheroid cell nuclei are labeled with Nunc blue (blue). 3D bi-photon imaging of spheroid infiltration by immune cells. **B** 10,000 CT26 cells deleted or not for MSH2 are growing for 3 days as spheroids and 100,000 PBMC from 3 spleens are activated with IL-15. Then, spheroids and PBMC are incubated for 3 days. PBMC were incubated before with anti-PD-1 (10 µg/mL). Flow cytometry staining of spheroids after dissociation. Quantification of differential immune checkpoint expression and lymphocyte infiltration according to the MSH2 status and ICB. N=6. Mann-Whitney U Test, p value<0.05 **C** Cytokine production in spheroids measured by Luminex (pg/mL). N=4. Mann-Whitney U Test, p value<0.05. **D** 10,000 B16F10 cells deleted or not for MSH2 are growing for 3 days as spheroids and 100,000 PBMC from 3 spleens are activated with IL-15. Then, spheroids and PBMC are incubated for 3 days. PBMC were incubated before with anti-PD-1 or anti-CTLA4 (10 µg/mL). Flow cytometry staining of spheroids after dissociation. Quantification of differential immune checkpoint expression and lymphocyte infiltration according to the MSH2 status and ICB. N=6. Mann-Whitney U Test, p value<0.05

**Extended Figure 13.**
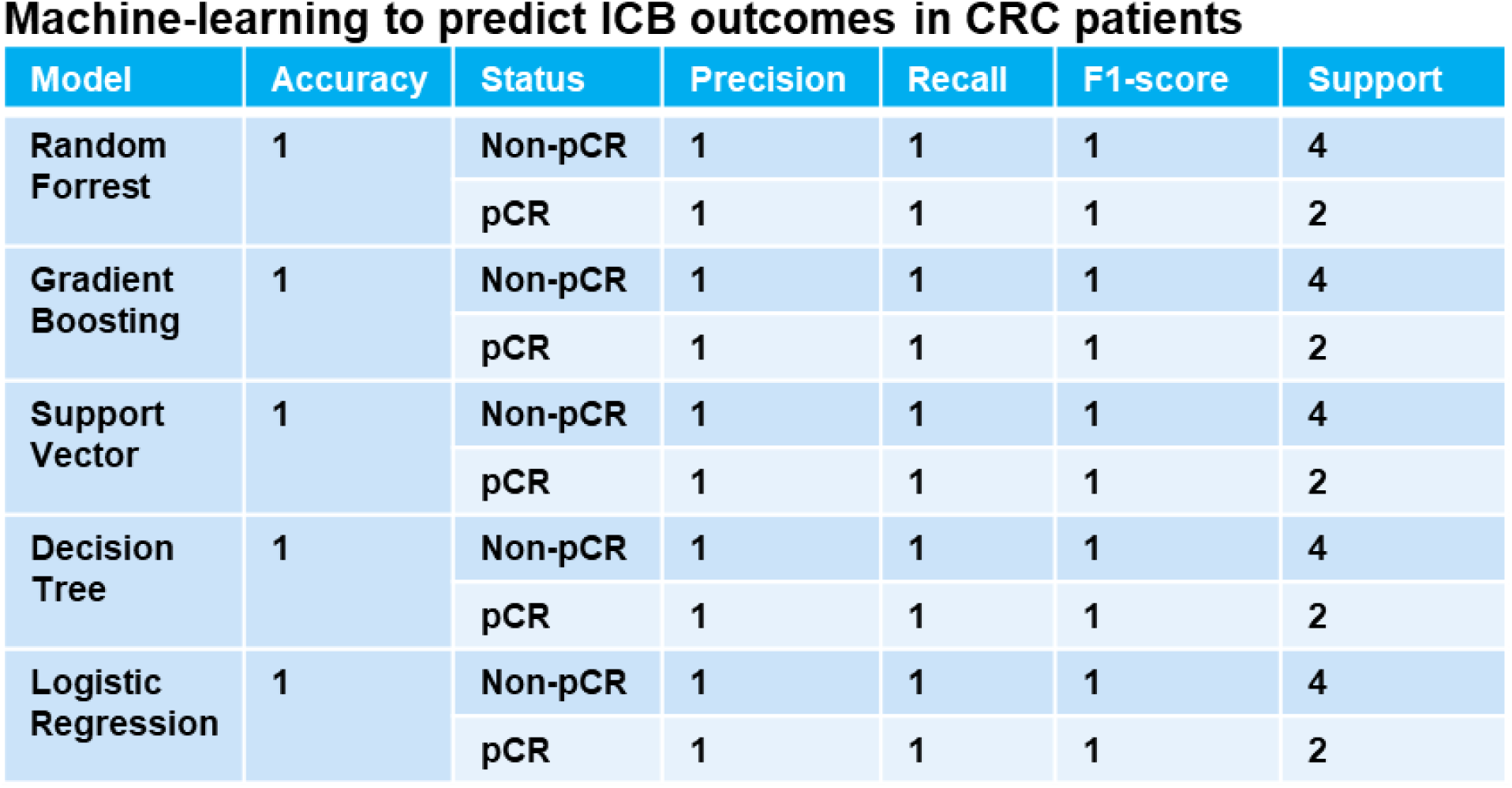
Machine-learning to predict ICB outcomes in CRC patients. Cell paper and KEYNOTE 177 Gastroenterology paper data : n = 29 patients ^13,34^. Responders n = 12. Non-responders n = 17. More A*03:01 in responders vs non-responders (Chi2 = 5.117844498910676, p-value = 0.023681014076523402). No statistical difference for other HLA. More MSIH Status in responders (Chi2 = 6.674496076839828, p-value = 0.00978021596680159). Based on MSI status and A*03:01 data, 5 machine-learning models managed to predict CRC patient response to ICB with an accuracy of 100%.

**Extended Figure 14.**
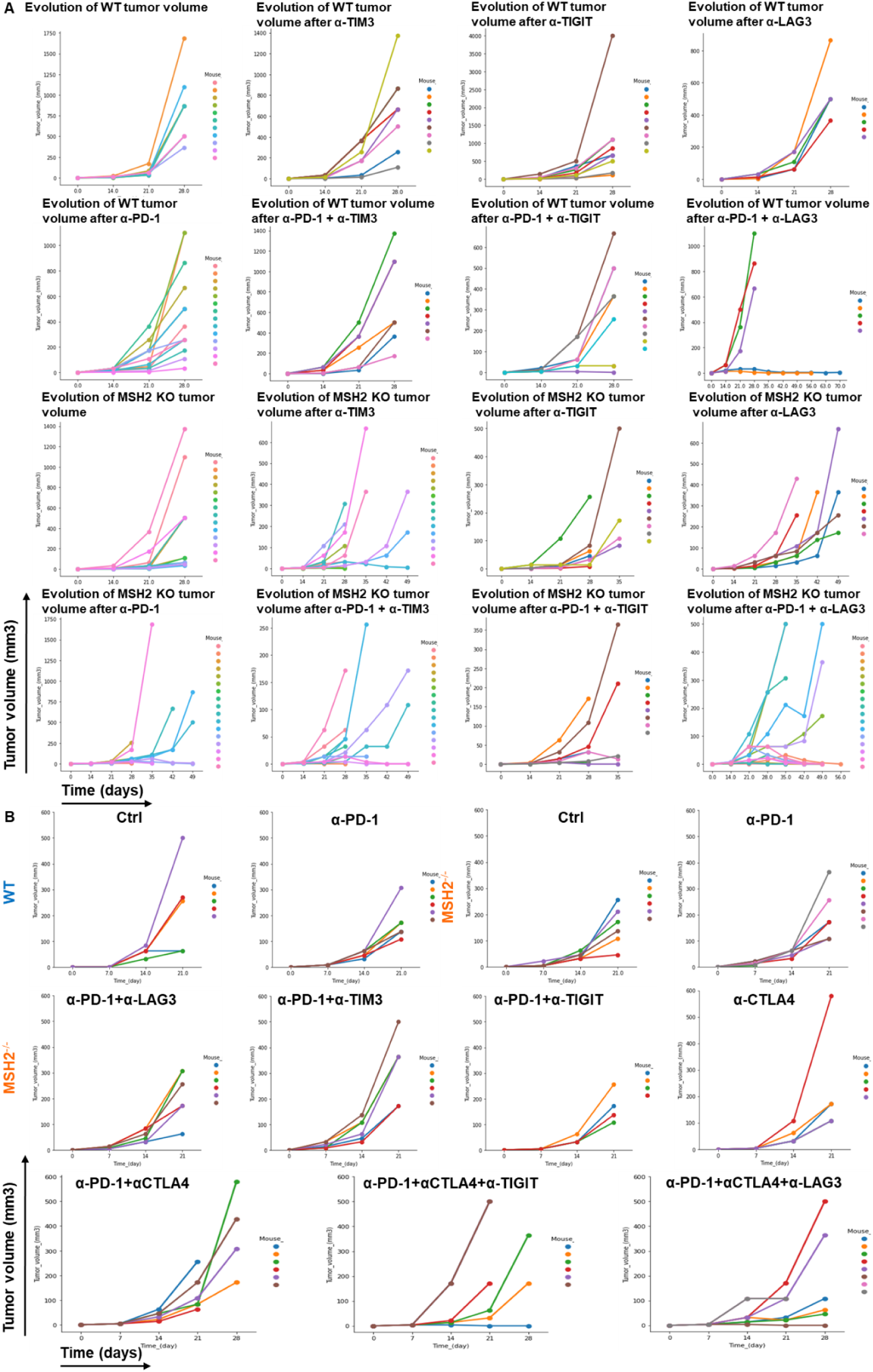
Impact of targeted multiple checkpoint combinations on MMRd tumor growth. Tumor volume in vivo over time of CT26 (**A**) and 4T1 (**B**) WT and MSH2 KO cells (200,000 cells/tumor) in the absence or presence of anti-PD-1, anti-TIM3, anti-LAG3, anti-TIGIT or anti CTLA4 (100 µg twice a week after 14 days) therapy (in mm3). N= 5 to 20 per arm. Mann-Whitney U Test, p value<0.05.

**Extended Figure 15.**
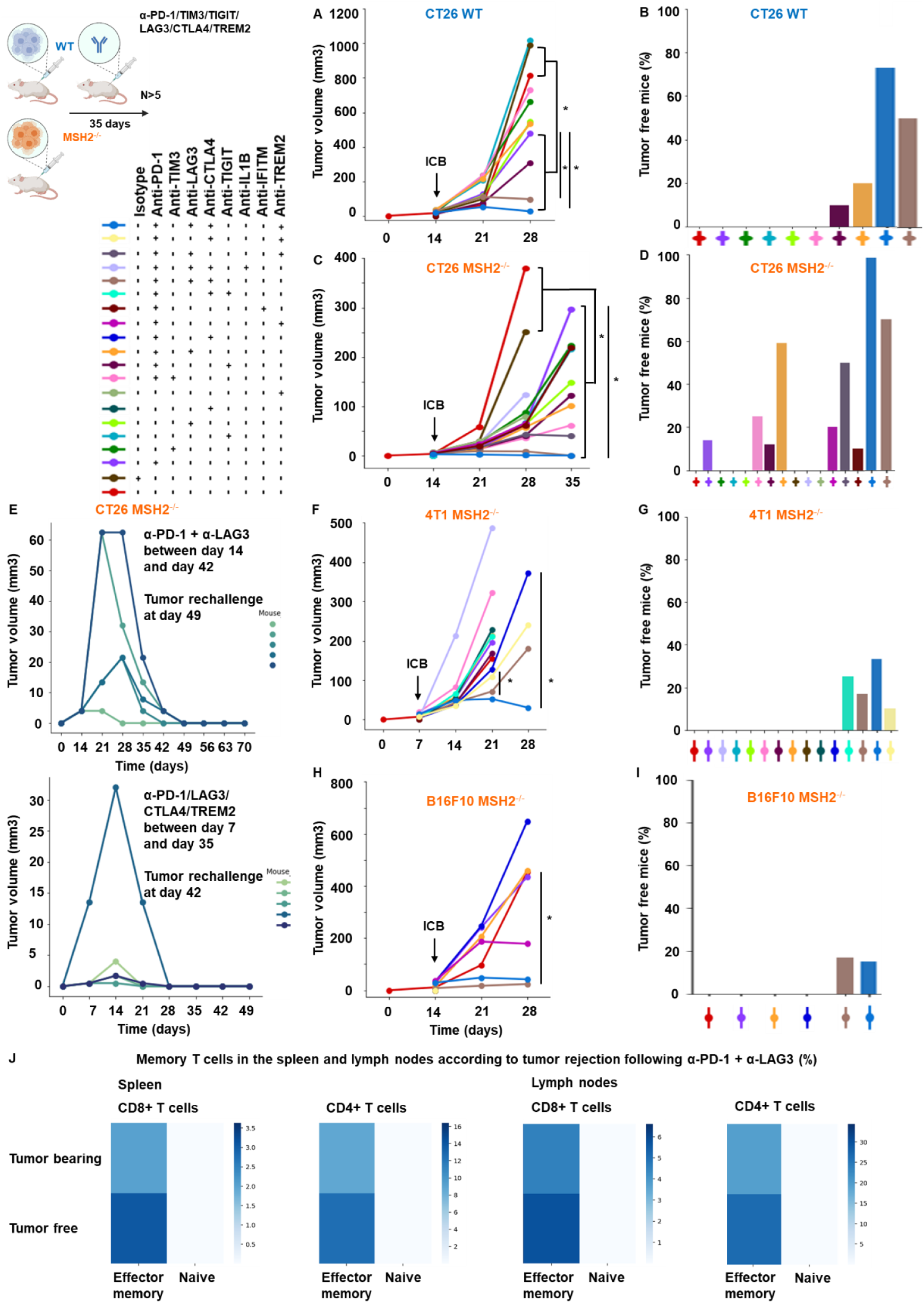
Myeloid and T cell targeting therapeutic combinations overcome resistance in both MMRd and MMRp tumors and elicit robust immune memory. A/C/F/H. Tumor volume (mean) in vivo over time of CT26, B16F10 and 4T1 WT and MSH2 KO cells (200,000 cells/tumor) in the absence or presence of anti-PD-1, anti-TIM3, anti-LAG3, anti-TIGIT, anti-IL1B, anti-TREM2, anti-IFITM or anti CTLA4 (100 µg twice a week after 14 days) therapy (in mm3). N= 5 to 20 per arm. Mann-Whitney U Test, p value<0.05. **B/D/G/I** Complete response of CT26, B16F10 and 4T1 WT and MSH2 KO tumors after ICB combinations (%). **E** After CT26 MSH2 KO tumor elimination following anti-PD-1 + anti-LAG3 therapy, treatment was stopped and a second tumor was inoculated on the opposite flank and its growth was measured for 3 weeks. N= 5. **J** Percentage of memory T cells in spleens and lymph nodes of mice that rejected or not a CT26 MSH2 KO tumor following anti-PD-1 + anti-LAG3 after 49 days (flow cytometry).

**Extended Figure 16.**
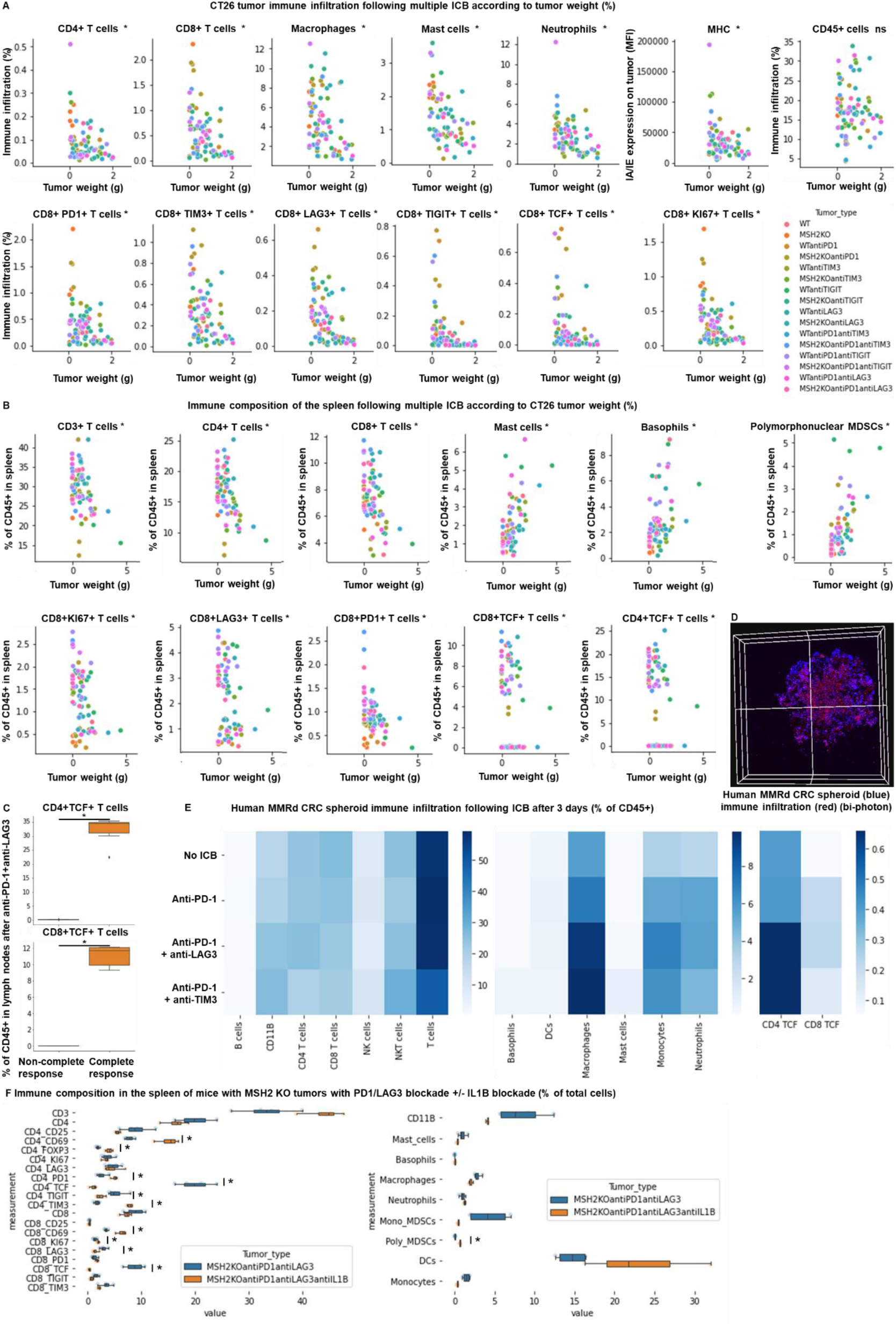
TCF+ T cells, neutrophils and macrophages orchestrate the response to targeted checkpoint/myeloid combinations. WT and MSH2 KO CT26 tumors are growing 28 days in absence or presence of anti-PD-1, anti-TIM3, anti-LAG3 or anti-TIGIT (100 µg twice a week after 14 days) therapy. **A** Linear regression, intratumoral immune infiltration and according to tumor weight, ICB and MSI status (flow cytometry). **B/C** Linear regression, immune composition in spleens and lymph nodes according to tumor weight, ICB and MSI status (flow cytometry). N= 79. Mann-Whitney U Test, p value<0.05. **D/E** 10,000 MMRd CRC patient-derived cells are growing for 3 days as spheroids and 100,000 PBMC from 3 donors are activated with IL-15. Then, spheroids and PBMC are incubated for 3 days. PBMC were incubated before with anti-PD-1, anti-TIM3 and anti-LAG3 (10 µg/mL). **D** Immune cells are previously labeled by cell tracker (red). Spheroid cell nuclei are labeled with Nunc blue (blue). 3D Bi-photon imaging. **E** Flow cytometry staining of spheroids after dissociation. Quantification of differential immune checkpoint expression and lymphocyte infiltration according to ICB. N=4. Mann-Whitney U Test, p value<0.05 **F** Immune composition in spleens of mice with MSH2 KO CT26 tumors according to the addition of PD1/LAG3 blockade +/-IL1B blockade (flow cytometry).

**Extended Figure 17.**
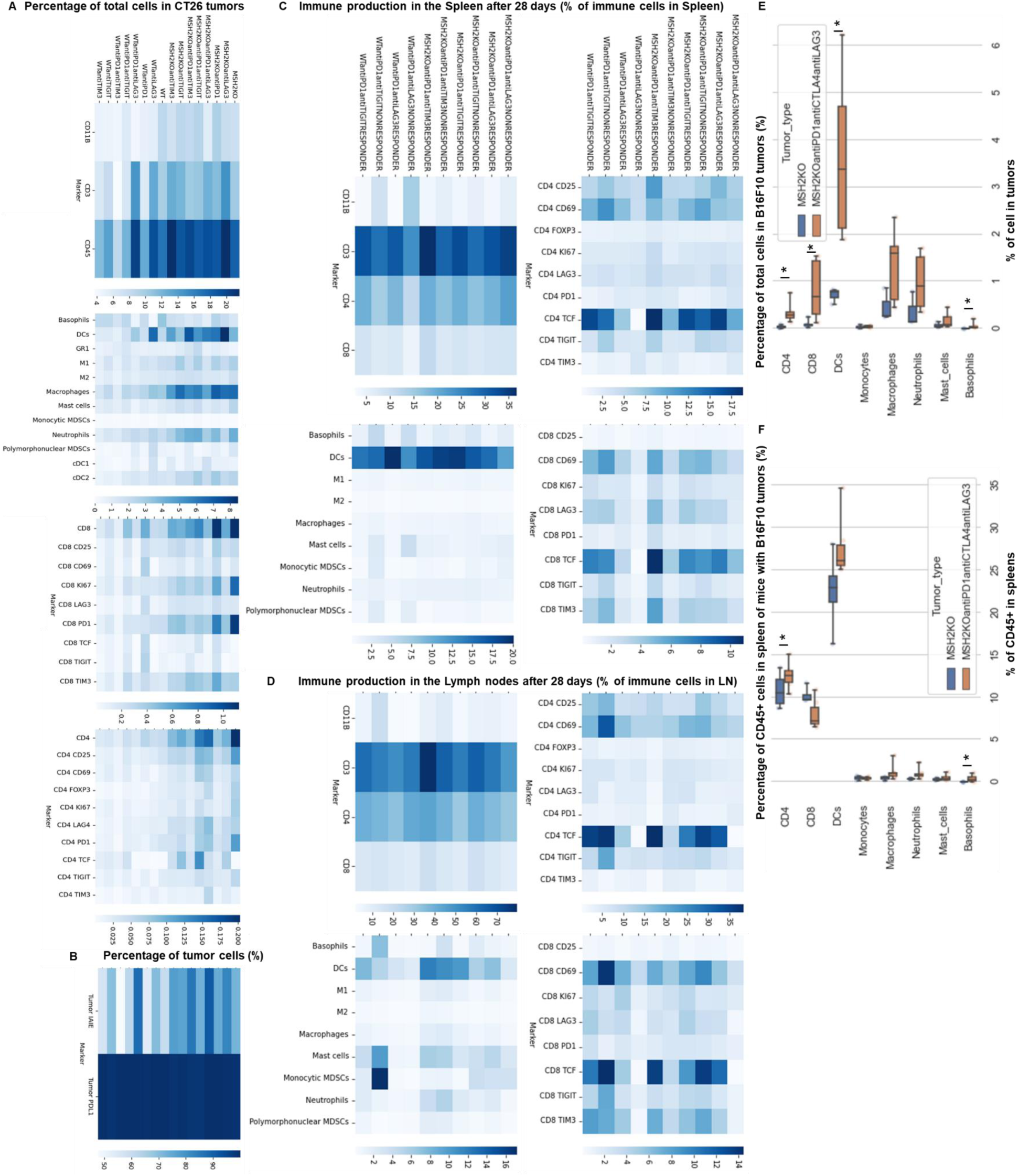
Impact of targeted multiple checkpoint combinations on MMRd tumor infiltration and immune composition in spleens and lymph nodes. CT26 and B16F10 WT and MSH2 KO tumors are growing 28 days in absence or presence of anti-PD-1, anti-TIM3, anti-LAG3 or anti-TIGIT (100 µg twice a week after 14 days) therapy. **A/B/E** Quantification by flow cytometry of differential immune checkpoint and lymphocyte infiltration expression according to the MSI status and ICB. **C/D/F** Immune composition in spleens and lymph nodes according to complete response following multiple ICB therapy. Responders are defined as free-tumor mice. N= 4. Mann-Whitney U Test, p value<0.05.

**Extended Figure 18.**
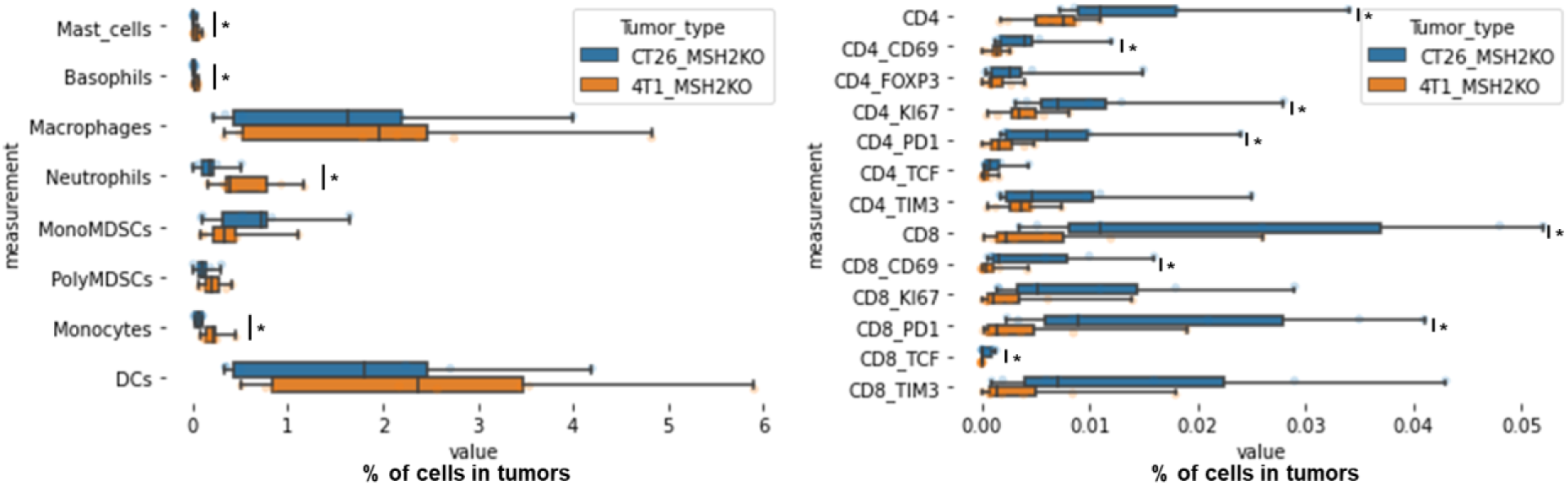
Immune infiltration of MSH2 KO CT26 and 4T1 tumors. CT26 and 4T1 tumors deleted or not for MSH2 are growing for 21 days in BALB/C mice. Quantification of differential immune checkpoint and lymphocyte infiltration expression according to the MSI status and ICB. N=8 replicates, Mann-Whitney U Test, p value<0.05

**Extended Figure 19.**
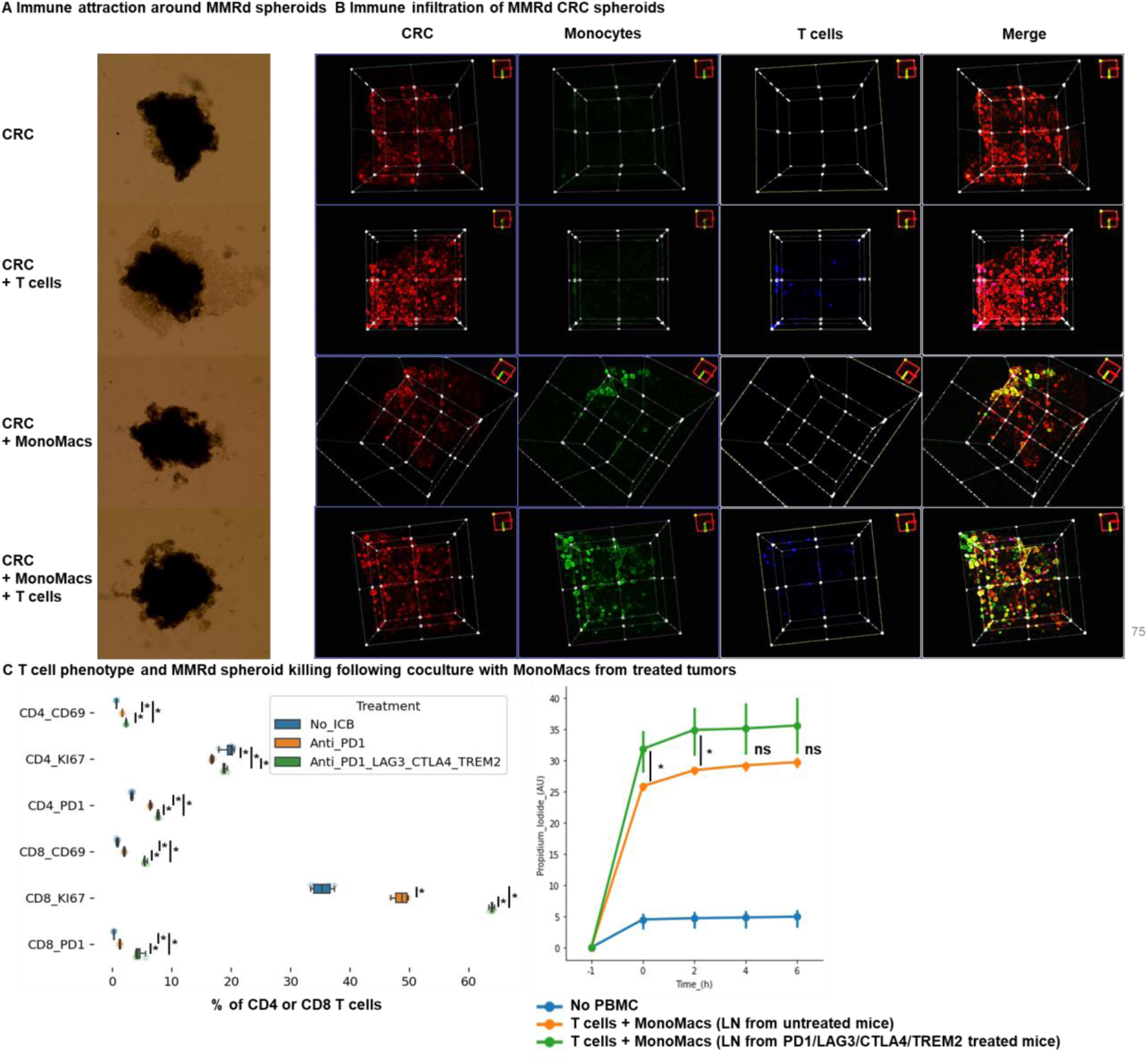
T cell and monocyte infiltration in human MMRd CRC spheroids. MMRd CRC human spheroids were amplified for 7 days, with n=10,000 CRC cells per spheroid and the Spherotribe kit (Idylle). In parallel, T cells and monocytes from blood were labeled with DeepRed/DeepGreen cell trackers for 30min respectively. Monocytes and T cells were added to spheroids (10,000 monocytes and 100,000 T cells per spheroid). **A** After 5 days, spheroid size and subsequent immune attraction was evaluated by microscopy and ImageJ. **B** Spheroid infiltration by monocytes and T cells and 3D aggregate formation were shown with BiPhoton. **C** Murine T cell phenotype and MMRd spheroid killing following coculture with MonoMacs from tumor or LN of mice bearing CT26 MSH2 KO tumors treated with anti-PD-1 or anti-PD-1/LAG3/CTLA4/TREM2. N= 4 replicates, Mann-Whitney U Test, p value<0.05

## Declaration of generative AI and AI-assisted technologies in the writing process

During the preparation of this work, the author(s) used ChatGPT and MistralAI in order to correct grammar and spelling issues. After using this tool/service, the author(s) reviewed and edited the content as needed and take full responsibility for the content of the publication.

